# A prefrontal network model operating near steady and oscillatory states links spike desynchronization and synaptic deficits in schizophrenia

**DOI:** 10.1101/2022.06.10.495666

**Authors:** David A. Crowe, Andrew Willow, Rachael K. Blackman, Adele L. DeNicola, Matthew V. Chafee, Bagrat Amirikian

## Abstract

Schizophrenia results in part from a failure of prefrontal networks but we lack full understanding of how disruptions at a synaptic level cause failures at the network level. This is a crucial gap in our understanding because it prevents us from discovering how genetic mutations and environmental risks that alter synaptic function cause prefrontal network to fail in schizophrenia. To address that question, we developed a recurrent spiking network model of prefrontal local circuits that can explain the link between NMDAR synaptic and spike timing deficits we recently observed in a pharmacological monkey model of prefrontal network failure in schizophrenia. We analyze how the balance between AMPA and NMDA components of recurrent excitation and GABA inhibition in the network influence spike timing to inform the biological data. We show that reducing recurrent NMDAR synaptic currents prevents the network from shifting from a steady to oscillatory state in response to extrinsic inputs such as might occur during behavior. This explains how NMDAR synaptic deficits, implicated by genetic evidence as causal in schizophrenia, could prevent the emergence of 0-lag synchronous spiking in prefrontal local circuits during behavior, potentially disconnecting those circuits via spike-timing dependent mechanisms in the human disease.

## Introduction

Schizophrenia, a devasting and still poorly understood disease (Insel, 2010; Jauhar et al., 2022), appears to result in part from the functional disruption of prefrontal cortical networks. We do not yet have a clear understanding of how or why this occurs, particularly at the cellular and synaptic level, where critical changes in cell and synaptic function lead to wide-scale failure of networks. To help fill this gap in knowledge, we have developed parallel nonhuman primate (Kummerfeld et al., 2020; Zick et al., 2018) and mouse genetic models (Zick et al., 2021) of prefrontal network failure in schizophrenia and used extracellular neural recording in these models to describe how insults associated with schizophrenia impact neural dynamics in prefrontal local circuits. We found that diverse insults operating through different molecular mechanisms (blocking NMDAR in primates or deleting a schizophrenia risk gene in mice related to miRNA processing) led to the same endpoint in prefrontal cortex: both perturbations reduced 0-lag synchronous spiking between prefrontal neurons along with a reduction in the amount of information transmitted by synaptic interactions between the neurons (Kummerfeld et al., 2020; Zick et al., 2021, 2018). This work suggests that multiple risk factors produce convergent effects on prefrontal networks during schizophrenia pathogenesis, but they do not address how or why molecular disruptions of cell function lead to the distortions of network dynamics that we have observed in monkey prefrontal cortex, and that in humans are likely responsible for the failure of prefrontal circuits and ultimately the cognitive deficits in the disease (Jauhar et al., 2022).

To discover how synaptic and network dynamical disturbances may be linked in schizophrenia, here we develop a spiking neural network model that was specifically informed by, and can provide mechanistic explanation for, neural recordings our group has obtained in monkey prefrontal cortex before and during NMDA receptor blockade (Zick et al., 2018). We use network stability and mean field analyses to investigate how the balance between NMDA and AMPA components of recurrent excitatory and GABA inhibitory currents influence regimes of network dynamics and spiking synchrony. For cortical neurons synchrony can occur naturally due to the local recurrent network connectivity, even when external afferent inputs are entirely uncorrelated. Theoretical studies have shown that such synchrony can arise in randomly connected recurrent networks operating in asynchronous irregular (Amit, 1989; Amit and Brunel, 1997; Brunel, 2000; Renart et al., 2010; van Vreeswijk and Sompolinsky, 1996; Vicente et al., 2008) and synchronous irregular regimes (Brunel, 2000; Brunel and Hakim, 1999; Brunel and Wang, 2003; Ledoux and Brunel, 2011). In both regimes individual neurons fire spikes highly irregularly at low rates, a typical situation in a cortex. The major distinction is that in an asynchronous regime population spike rate is steady in time, whereas in a synchronous regime it becomes oscillatory. We show that simulated prefrontal networks operating near the boundary between steady (asynchronous irregular) and oscillatory (synchronous irregular) regimes in the synaptic parameter space can explain several key experimental observations. First, such networks achieve biologically realistic stochastic spike trains and firing rates of excitatory and inhibitory neurons in prefrontal cortex. Second, increased extrinsic inputs such as might occur during behavior shift these networks from a steady into an oscillatory regime that causes the emergence of 0-lag spiking between neurons as they stochastically entrain to oscillatory population activity. Third, and perhaps most importantly, we show that reducing recurrent NMDAR synaptic currents prevents these networks from transitioning into oscillatory activity in response to extrinsic inputs, thereby preventing the emergence of 0-lag spike synchrony. This allows us to establish strong parallels between simulated and biological data, including the emergence of 0-lag synchronous spiking via recurrent synaptic interactions between neurons during behavior, the association between synchronous spiking and oscillatory population activity, as well as their joint dependence on NMDAR synaptic mechanisms, both in our current simulation and in the neural data (Zick et al., 2018).

These theoretical results suggests that NMDAR synaptic mechanisms control oscillatory population activity and synchronous spiking between prefrontal neurons. The impact of NMDAR synaptic transmission modulation on neural dynamics have been simulated in artificial neural networks in various contexts; for instance, stability of working memory (Compte et al., 2000; Wang, 1999), oscillatory activity observed in LFP recordings (Brunel and Wang, 2003), or EEG correlates of entrainment of auditory cortex neurons to periodic auditory stimuli (Kirli et al., 2014). However, NMDAR conductance modulation in these simulations was carried out for networks in which either the very nature of simulated spiking dynamics was incompatible with the spiking dynamics observed in prefrontal neurons (Kirli et al., 2014), or the effect of NMDAR modulation did not have impact on spiking synchrony and oscillations (Brunel and Wang, 2003), or it was opposite to the effect in the question (Compte et al., 2000) that we observed in prefrontal cortex recordings during a behavioral task measuring specific deficits in schizophrenia (Zick et al., 2018) and that we capture in the current network model. Thus, no previous modeling studies have addressed the issue of how NMDAR hypofunction can desynchronize spiking activity in prefrontal cortex, nor how spiking synchrony is modulated during transitions between oscillatory and steady states, or how such transitions may be influenced by NMDAR synaptic mechanisms. By presenting evidence that prefrontal networks operate near the boundary between steady (asynchronous irregular) and oscillatory (synchronous irregular) regimes, the current study provides a key conceptual advance and unique circuit level insight explaining why deficits in NMDAR synaptic transmission (such as are thought to occur in schizophrenia) reduce oscillations in prefrontal networks and synchronous spiking between prefrontal neurons, as it is observed in neurophysiological experiments modeling prefrontal network failure in schizophrenia (Zick et al., 2018).

The interdependence between NMDAR synaptic mechanisms, oscillatory dynamics and spike timing in prefrontal local circuits may be of critical importance to understanding the etiopathogenesis of schizophrenia. Synchronous, or near synchronous spiking in pre- and postsynaptic neurons governs spike timing dependent synaptic plasticity (Dan and Poo, 2004). Thus, any insult or perturbation (either of genetic, developmental, or environmental origin) that disrupts the balance between NMDAR synaptic inputs, oscillatory activity and spike timing in prefrontal local circuits could ultimately cause these circuits to actively disconnect via spike-timing dependent synaptic plasticity, contributing to disease progression (Zick et al., 2021, 2018). Additionally, recent evidence suggests that synchronous pre- and postsynaptic spiking establishes an eligibility trace at specific dendritic spines rendering them susceptible to potentiation by subsequent dopamine (He et al., 2015; Kasai et al., 2021; Yagishita et al., 2014). This suggests that network oscillations, and the spike synchrony they create, could confer synapse specificity on dopamine signals, effectively linking Hebbian synaptic plasticity and reinforcement learning mechanisms in the brain. Therefore, understanding the synaptic mechanisms that underlie network oscillations and synchronous spiking in recurrent prefrontal networks may help us understand how spike timing may govern reinforcement learning in these networks normally, and how NMDAR synaptic deficits, implicated as causal in schizophrenia, may disrupt learning and functional integrity of these networks in disease.

## Results

### Summary of experimental results

In this section we summarize main experimental findings reported previously by our group (Zick et al., 2018). In that study, spike trains of ensembles of single neurons were recorded simultaneously from PFC of monkeys while they performed the dot-pattern expectancy (DPX) task, a task that measures specific deficits in cognitive control in schizophrenia (Jones et al., 2010). In the DPX task, the correct response (left or right joystick movement) to a probe stimulus depends on a preceding cue followed by a delay period (Methods).

In the present study, we focus on PFC population spike dynamics recorded in the DPX task under two conditions: drug-naïve and drug. The drug naïve data were collected before monkeys were administered drug, phencyclidine, which is an NMDA receptor antagonist. Figure 1 shows the population average pairwise correlation between spike trains of neurons recorded in drug-naïve (black) and drug (magenta) conditions. The strength of spike correlation was quantified by the ratio between the observed frequency of synchronous spikes (2ms resolution) and the frequency expected if the spike trains were uncorrelated (we subtracted 1 from this ratio so that correlation value is zero for uncorrelated, positive for correlated, and negative for anticorrelated spike activity, Methods). The frequency of spike synchrony was determined from activity observed during a short (100 ms-long) window that was slid across time of task performance. Figure 1a shows that spike synchrony obtained from trials aligned to the cue onset (time 0) remained relatively weak and unchanged during the cue and delay periods, until the probe onset, in both drug-naïve and drug conditions. Because the instant of response after probe presentation varied from trial to trial, to appreciate the time course of synchrony after delay period immediately preceding the response, in Fig.1b we aligned trials to response time (time 0). It is seen that spike synchrony started to increase sharply about 200ms before the motor response in the drug-naïve condition. In contrast, it practically did not change in the drug condition. We term this effect as NMDAR blockage induced desynchronization of spiking activity.

**Figure 1.**
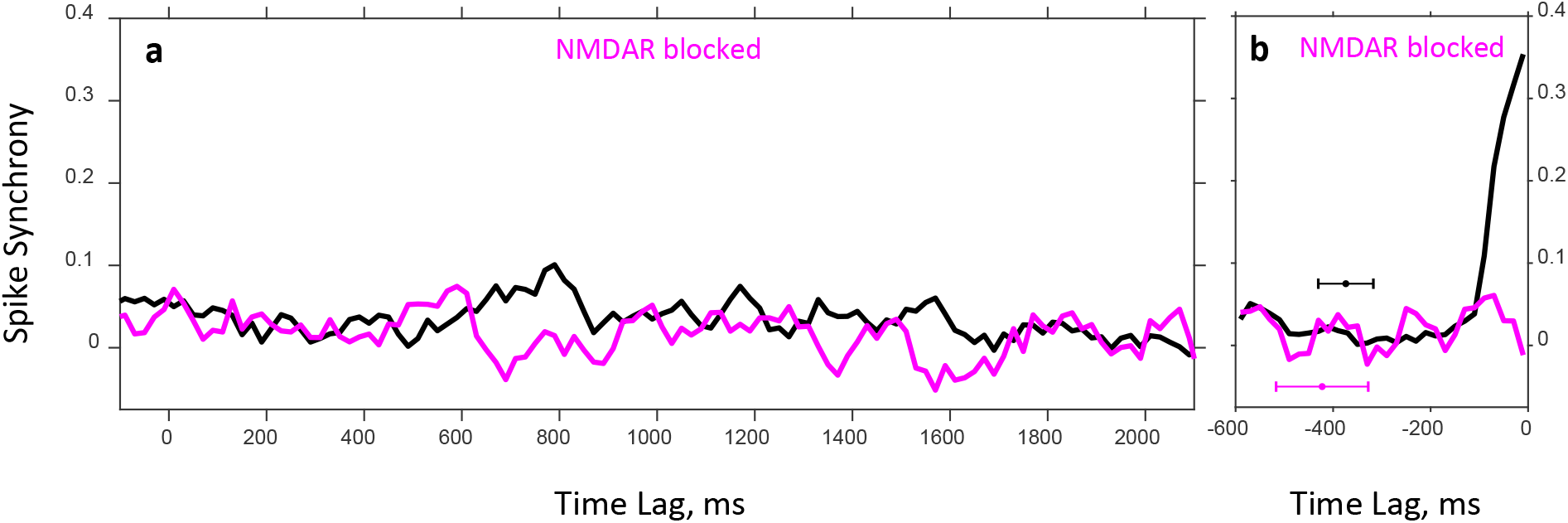
Population average spike synchrony between spike trains of neuron pairs recorded during the DPX task as a function of time. Plots are shown for drug-naïve (black) and drug (magenta) conditions. Spike synchrony was measured with 2 ms resolution, and its temporal modulation was estimated with 100 ms resolution. Only neuron pairs for which a reasonably reliable estimation of synchrony could be achieved contributed to the plots (see Methods). **a**: Trials are aligned to the cue onset (*t* = 0 ms); in all trials, the cue was presented until *t* = 1,000 ms, followed by a 1,000 ms delay period, after which the probe was presented at *t* = 2,000 ms. The numbers of contributing neuron pairs for drug-naïve and drug conditions are 978 and 368, respectively. **b**: Trials are aligned to the time of motor response (*t* = 0 ms) to show the temporal modulation of synchrony during the last 600 ms immediately preceding the response. Color-coded horizontal error-bars indicate the mean and standard deviation of the probe presentation time for the corresponding drug condition. The numbers of contributing neuron pairs for drug-naïve and drug conditions are 1,246 and 434, respectively.

### Network model and theoretical framework

To understand the phenomenon of drug-induced desynchronization of spiking activity and the role played by various components of synaptic currents, we considered a spiking network model representing a local circuit of monkey PFC. Details of the model and the theoretical framework are given in Methods. Here, we only highlight their main aspects.

The network comprises excitatory and inhibitory neurons representing populations of pyramidal cells and interneurons, respectively. Synaptic connections are random and sparse, but the number of connections received by individual neurons is large. In addition to the recurrent local connections, each neuron also receives external connections from excitatory neurons outside of the network that fire spikes with rate *ν*_X_.

Recurrent synaptic currents of excitatory connections are two-component, mediated by AMPA and NMDA receptors, whereas currents of inhibitory connections are mediated by GABA receptors. External currents represent the noisy inputs due to the background synaptic activity and mediated by AMPA receptors. Thus, the model entails eight maximal synaptic conductance parameters *g*_X,*α*_, *g*_AMPA,*α*_, *g*_NMDA,*α*_, *g*_GABA,*α*_ corresponding to the external AMPA, recurrent AMPA, NMDA, and GABA currents (*α* = E,I for excitatory and inhibitory neurons, respectively).

To produce a desired regime of network dynamics (asynchronous or synchronous) with a given firing rate of excitatory and inhibitory neurons *ν*_E_ and *ν*_I_, respectively, the values of the conductance parameters should be properly adjusted. For this purpose, we used mean field analysis. In this framework, population mean firing rates 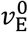 and 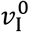 in the asynchronous stationary state of the network can be effectively parametrized by three parameters expressed as ratios of component synaptic currents: *I*_AMPA_/*I*_GABA_, *I*_NMDA_/*I*_GABA_, and *I*_X,E_/*I*_*θ*,E_, where*I_R_* is the mean current of the *R*-receptor mediated synapse (*R* = X, AMPA, NMDA, GABA), and *I*_*θ*,E_ is the current that is needed for an excitatory neuron to reach firing threshold *θ* in absence of recurrent feedback. These parameters characterize the balance between recurrent excitation and inhibition, and the balance between external input and firing threshold. Once they are specified, for a given external spike rate *ν*_X_ one can solve the mean field equations to obtain the underlying eight synaptic conductances providing the desired population mean firing rates 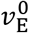 and 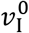 in asynchronous state of the network.

While the mean field analysis allows us to determine synaptic conductances that achieve desired firing rates of neurons, whether these rates remain stable over time is another issue. To address it, we conduct a linear stability analysis of the asynchronous state to understand if the network develops oscillatory instability caused by small fluctuations in population firing rates. This analysis entails two parameters, *λ* and ω, describing the rate of instability growth and the oscillation frequency. The asynchronous state is stable when *λ* < 0; in this case small perturbations of firing rates cause exponentially damped oscillation of network activity. The case *λ* = 0 corresponds to the onset of instability of the asynchronous state and the emergence of sustained sinusoidal oscillations of population average firing rates with frequency ω; in the oscillatory regime spike trains remain sparse and irregular but at each oscillation cycle a random subset of network neurons fire synchronously giving rise to the synchronous irregular state. Lastly, when *λ* >0, small fluctuations in the stationary rates develop oscillatory instability with the amplitude of oscillations growing exponentially in time; however, higher order terms neglected in linear analysis can eventually saturate the instability growth (Brunel and Hakim, 1999), resulting in a stable oscillation with a finite amplitude.

To examine the boundary between the regions of asynchronous and synchronous states, we fix the balance of tonic NMDA current relative to GABA current, *I*_NMDA_/*I*_GABA_, and vary the remaining two parameters: the balance between recurrent excitation and inhibition, *I*_AMPA_/*I*_GABA_, and the balance between external excitation and firing threshold, *I*_X,E_/*I*_θ,E_. For a given point in this (*I*_AMPA_/*I*_GABA_, *I*_X,E_/*I*_*θ*,E_) parameter plane we solve the mean field equations to find the underlying set of eight synaptic conductances that provide the prescribed rates 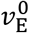 and 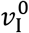 given external spike rate *ν*_X_, and then carry out linear stability analysis to find out if these rates are stable. Figure 2a shows a state diagram of the system for which external spike rate is set to *ν*_X_ = 5 Hz, the rates of excitatory and inhibitory populations are set to 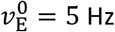, 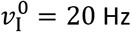, and the NMDA current balance is fixed at *I*_NMDA_/*I*_GABA_ = 0.2. The diagram shows solutions for *λ* obtained from the linear stability analysis in the (*I*_AMPA_/*I*_GABA_, *I*_X,E_/*I*_*θ*,E_) parameter space. The asynchronous stationary state corresponds to the region where *λ* < 0, whereas the synchronous oscillation state is realized in the region where *λ* >0. The asynchronous and synchronous states are separated by a “critical” or instability line on which *λ* = 0 (shown in white color in Fig. 2a). This boundary is the locus where the stationary network dynamics becomes unstable, and the sinusoidal oscillation of network activity develops. The oscillation frequency on the critical line, *f*_ntwrk_ = *ω*/2*π*, as a function of the balance between the recurrent AMPA and GABA currents, *I*_AMPA_/*I*_GABA_, is shown in Fig. 2b.

**Figure 2.**
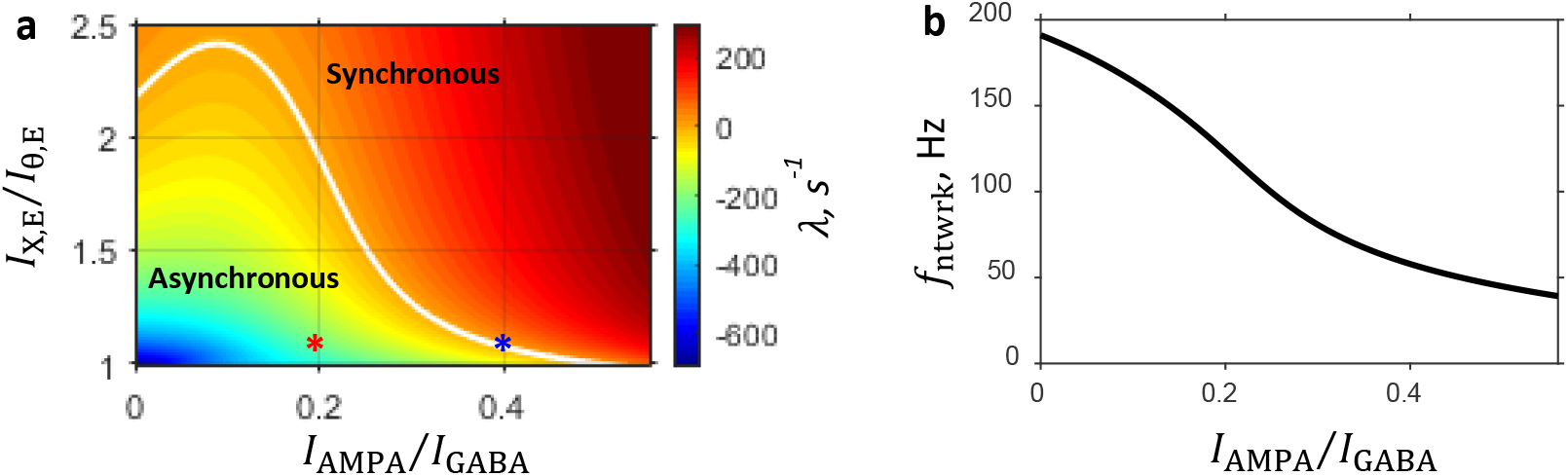
Characteristics of the system predicted by the linear stability analysis. Parameters are as follows: prescribed firing rates of excitatory and inhibitory populations are 5 Hz and 20 Hz, respectively; external input spike rate is 5 Hz; and the balance between NMDA and GABA currents is fixed at 0.2. **a:** State diagram in the (*I*_AMPA_/*I*_GABA_, *I*_X,E_/*I*_*θ*,E_) parameter plane showing color coded value of the rate of instability growth *λ*: in the region of the parameter space where *λ* <0 the asynchronous state is stable, whereas the region where *λ* >0 corresponds to the synchronous oscillation state. The two regimes are separated by a critical line on which *λ* = 0. This boundary, shown by a white line, is the locus where the stationary network dynamic becomes unstable, and oscillatory population activity develops. Each point in this parameter plane corresponds to a network with a specific set of eight synaptic conductances provided by the mean field approximation. Red and blue asterisks are the points in the state diagram corresponding to the steady and critical primary networks, respectively (see Selection of Primary Networks in Results). **b:** Network oscillation frequency that develops on the critical line as a function of the balance between AMPA component of recurrent excitation and inhibition. **Figure supplement 1.** Dependence of the characteristic features of the network on the balance between the NMDA and GABA currents.

The characteristic features of the state diagram qualitatively remain unchanged when the balance between the NMDA and GABA currents is varied (Fig. 2–figure supplement 1a). Furthermore, the network frequency at the onset of oscillation, *f*_ntwrk_, essentially is independent of the *I*_NMDA_/*I*_GABA_ balance (Fig. 2–figure supplement 1b).

### Integration of DPX task context and drug condition into the model

To study spike synchrony in asynchronous and synchronous networks in the context of the DPX task performed in drug-naïve and drug conditions (Zick et al., 2018), we make two assumptions regarding neural and synaptic activity: 1) the increase in spike synchronization observed before the monkey’s response in (Zick et al., 2018) is due to task-specific external afferent signals received by PFC neurons after probe presentation; 2) administration of NMDAR antagonist results in blocking NMDAR mediated synaptic currents. In the framework of our model, we implemented these assumptions as follows: task specific external signals were accounted for by an increase in the external spike rate from its background level *ν*_X_, whereas the effect of drug administration was modeled by setting NMDAR conductances *g*_NMDA,E_ and *g*_NMDA,I_ to zero.

Next, to investigate how spike synchrony in asynchronous and synchronous networks depends on the modulations of *ν*_X_ and *g*_NMDA,α_, for each network regime we proceed with the following three steps. First, we choose proper values for conductances, so that the underlying network operates in a desired regime providing the prescribed population firing rates 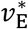 and 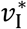 for a given external spike rate 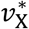. We shall designate this network as the *primary network* relating to the underlying regime and distinguish the corresponding values of all its parameters by the asterisk (*). Second, we carry out a series of network simulations, in which external spike rate *ν*_X_ and NMDAR conductance *g*_NMDA,α_ are varied relative to their standard values 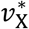 and 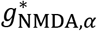, respectively. Lastly, for each simulated network, we compute population average pairwise correlation between spike trains of neurons and analyze how this correlation depends on the external spike rate and NMDAR conductance.

### Selection of primary networks

To perform a comparison between the primary networks, we need to choose appropriate values for their parameters. We begin with the parameters that are common to both networks. First, we set the excitatory and inhibitory population mean firing rates to 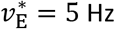 and 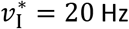, respectively, which are on the order of magnitude of spontaneous rates observed for PFC neurons. Second, since external inputs represent activity of excitatory neurons outside the PFC circuit model, we choose the background external rate 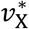 to be the same as the excitatory population rate 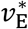 inside the model and, thus, set 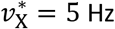. Lastly, for both networks we fix the balance between NMDA and GABA currents at 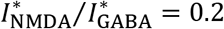. Note that the state diagram in the (*I*_AMPA_/*I*_GABA_, *I*_X,E_/*I*_*θ*,E_) space shown in Fig. 2a was obtained exactly for these values of the above listed parameters. We use this state diagram for selecting the primary networks and determining the remaining parameters that are network specific.

In this regard, we note that each point in the (*I*_AMPA_/*I*_GABA_, *I*_X,E_/*I*_*θ*,E_) plane corresponds to a network with a specific set of synaptic conductances. For synchronous regime, we look for a network on the critical line (*λ* = 0, white line in Fig. 2a), at the onset of oscillatory instability with a frequency in the *γ*-band (a frequency band associated with the LFPs recorded from prefrontal areas (Bastos et al., 2018; Lundqvist et al., 2016)). For instance, the point marked by a blue asterisk in Fig. 2a located at 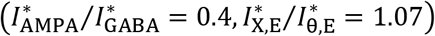 corresponds to such a network with oscillation frequency 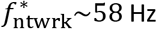 (cf. Fig. 2b). In the following, we refer to this network as the *critical state primary network*.

Correspondingly, for the asynchronous regime, we need to select a network that is far from the critical line and deep in the region of stable network dynamics (*λ* < 0). The point marked by a red asterisk in Fig. 2a located at 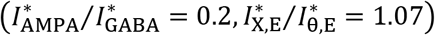 is an example of such a network. We shall refer to this network as the *steady state primary network*. For each primary network, we obtain the underlying set of eight synaptic conductance parameters 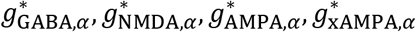 (*α* = E,I) by numerically solving the mean field equations.

### Correlation of spiking activity and synchrony in the asynchronous and synchronous states

To investigate characteristic features of spiking dynamics in asynchronous and synchronous regimes, we carried out direct simulations of the primary networks. Both networks comprise *N* = 5,000 neurons, of which *N*_E_ = 4,000 are excitatory and *N*_I_ = 1,000 inhibitory. Neurons are connected randomly with a probability *p* = 0.2. Figure 3 illustrate the behavior of simulated networks with synaptic conductance parameters corresponding to the steady and critical primary networks indicated by the red and blue asterisks, respectively, in the state diagram presented in Fig. 2a. The dynamic behavior is shown at the level of individual cell activity (spike rasters, top of panels in Fig.3), as well as whole population activity (bottom of panels in Fig.3).

**Figure 3.**
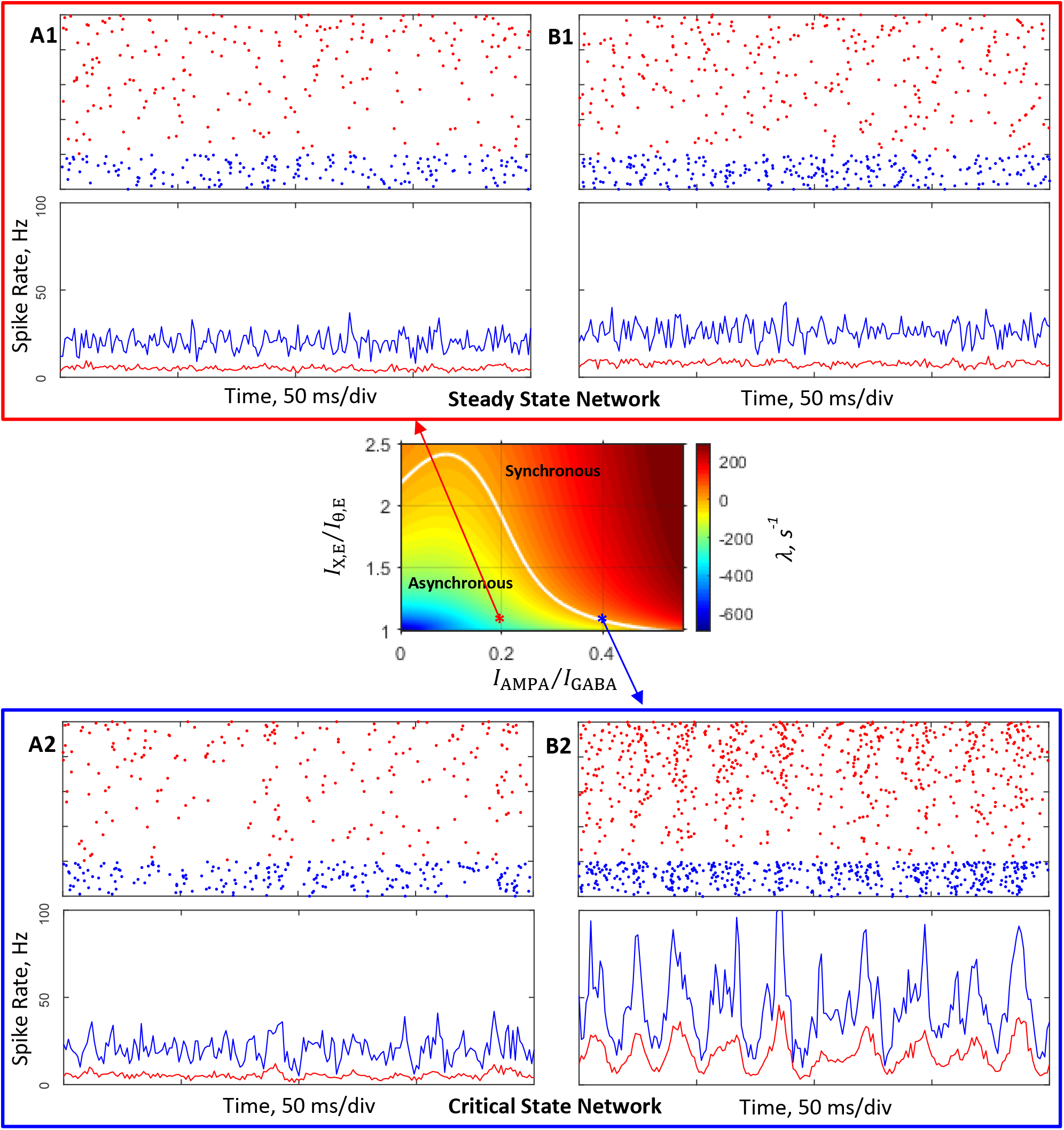
Simulations of networks composed of 4,000 excitatory and 1,000 inhibitory neurons connected randomly with probability 0.2. Conductance parameters are solutions of mean field equations for the steady state primary network (**a1**, **b1**) and the critical state primary network (**a2**, **b2**) corresponding to the red and blue asterisks, respectively, in the (*I*_AMPA_/*I*_GABA_, *I*_X,E_/*I*_*θ*,E_) state plane shown in Fig 2a and inset. **a1**, **b1**, **a2**, **b2**: Top, spike rasters (sorted by rate) of 200 excitatory (red) and 50 inhibitory (blue) neurons. Bottom, time-varying activity (1ms resolution) of excitatory (red) and inhibitory (blue) populations. **a1**, **a2**: External input spike rate *ν*_X_ = 5 Hz. Excitatory and inhibitory neurons display average firing rates of, respectively, 5.2 Hz and 20 Hz (**a1**), and 5.5 Hz and 21 Hz (**a2**). **b1**, **b2**: In these simulations *ν*_X_ was increased by 5%. Excitatory and inhibitory neurons display average firing rates of, respectively, 7.8 Hz and 26 Hz (**b1**), and 15 Hz and 42 Hz (**b2**). **Figure supplement 1.** Power spectra of population spiking activity observed in network simulations.

In simulations shown in Fig. 3 panels a1 and a2 external spike rate *ν*_X_ was fixed at the level of 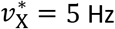 chosen for the primary networks. It is seen that excitatory and inhibitory neurons exhibit highly irregular firing with average rates, *ν*_E_ and *ν*_I_, about 5.2 Hz and 20 Hz in the steady state primary network (Fig. 3a1) and 5.5 Hz and 21 Hz in the critical state primary network (Fig. 3a2). These observed in simulations rates *ν*_E_ and *ν*_I_ are in good agreement with the prescribed rates 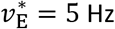 and 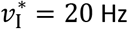 that were used to derive the synaptic conductance parameters of the simulated networks. Moreover, Fig. 3a1 demonstrates that population activity of the steady state primary network is rather stationary in time, whereas activity of the critical primary network shown in Fig. 3a2 exhibits signs of developing of oscillatory instability (compare Fig.3–figure supplement 1a1 vs Fig.3–figure supplement 1a2). Thus, spiking dynamics observed in the simulated steady state primary network displays basic characteristics of the asynchronous regime—irregular firing of individual neurons and stationary population activity. Correspondingly, the behavior of the simulated critical state primary network exhibits similarity with the boundary regime on which the asynchronous stationary state destabilizes and oscillatory behavior of the population activity emerges.

Panels b1 and b2 in Fig. 3 demonstrate results of simulations in which external spike rate *ν*_X_ was increased by 5% relative to the rate 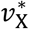 used in simulations illustrated in Fig. 3 panels a1 and a2. For the steady state primary network (Fig. 3b1), the firing rates of excitatory and inhibitory neurons increase with the external drive. However, the regime of network dynamics qualitatively does not change and remains asynchronous (compare Fig.3–figure supplement 1a1 vs Fig.3–figure supplement 1b1). In contrast, stronger external inputs received by the critical state primary network synchronize population activity (Fig. 3b2). It is seen that while individual neurons continue to fire irregularly, population activity now clearly exhibits oscillatory behavior, indicating that the network is in synchronous irregular regime in which the average firing frequency of neurons is low, about 20 Hz, compared to the frequency of network oscillation, which is about 50 Hz (see Fig.3–figure supplement 1b2). This frequency is close to the theoretically predicted network frequency of 58 Hz near the onset of oscillation.

Thus, direct simulations confirm that analytically derived network parameters for both steady and critical primary networks provide the anticipated regimes of network dynamics.

To facilitate the comparison of characteristic features exhibited by a simulated network with experimentally measurable quantities, we compute temporal correlation of spiking activity that quantifies average pairwise correlation between spike trains of excitatory neurons. In the context of the DPX task performed in drug-naïve and drug conditions studied in (Zick et al., 2018) and with the purpose of elucidating the mechanism of drug-induced desynchronization of spiking activity, we investigated how temporal correlations depend on the strength of external drive and the NMDAR mediated synaptic current. To this end, we varied external input rate *ν*_X_ and the NMDAR conductance parameters *g*_NMDA,E_ and *g*_NMDA,I_ relative to their respective standard values 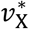, and 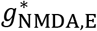 and 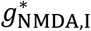, while keeping all other system parameters fixed, and performed simulations of the ensuing networks. Conductances for excitatory and inhibitory neurons were scaled with the same factor and, therefore, their relative values 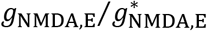 and 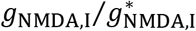 are the same; in the following we drop the E,I designation.

Figure 4 displays correlation of spiking activity (panels a1, a2, c1, c2) and synchrony (0-lag correlation, panels b1, b2, d1, d2) obtained from spike trains of simulated steady (panels a1, b1, c1, d1) and critical (panels a2, b2, c2, d2) networks for a range of 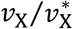 (panels a1, a2, b1, b2) and 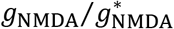 (panels c1, c2, d1, d2) values. It is seen that in the steady state primary network correlations are weak and insensitive to the modulations of external input rate or NMDAR conductance (Fig. 4 panels a1, b1, c1, d1). In contrast, in the critical state primary network temporal correlations show sharp dependence on these parameters (Fig. 4 panels a2, c2), and with increasing external drive or NMDAR conductance spike synchrony could be modulated from relatively weak to rather strong (Fig. 4 panels b2, d2).

**Figure 4.**
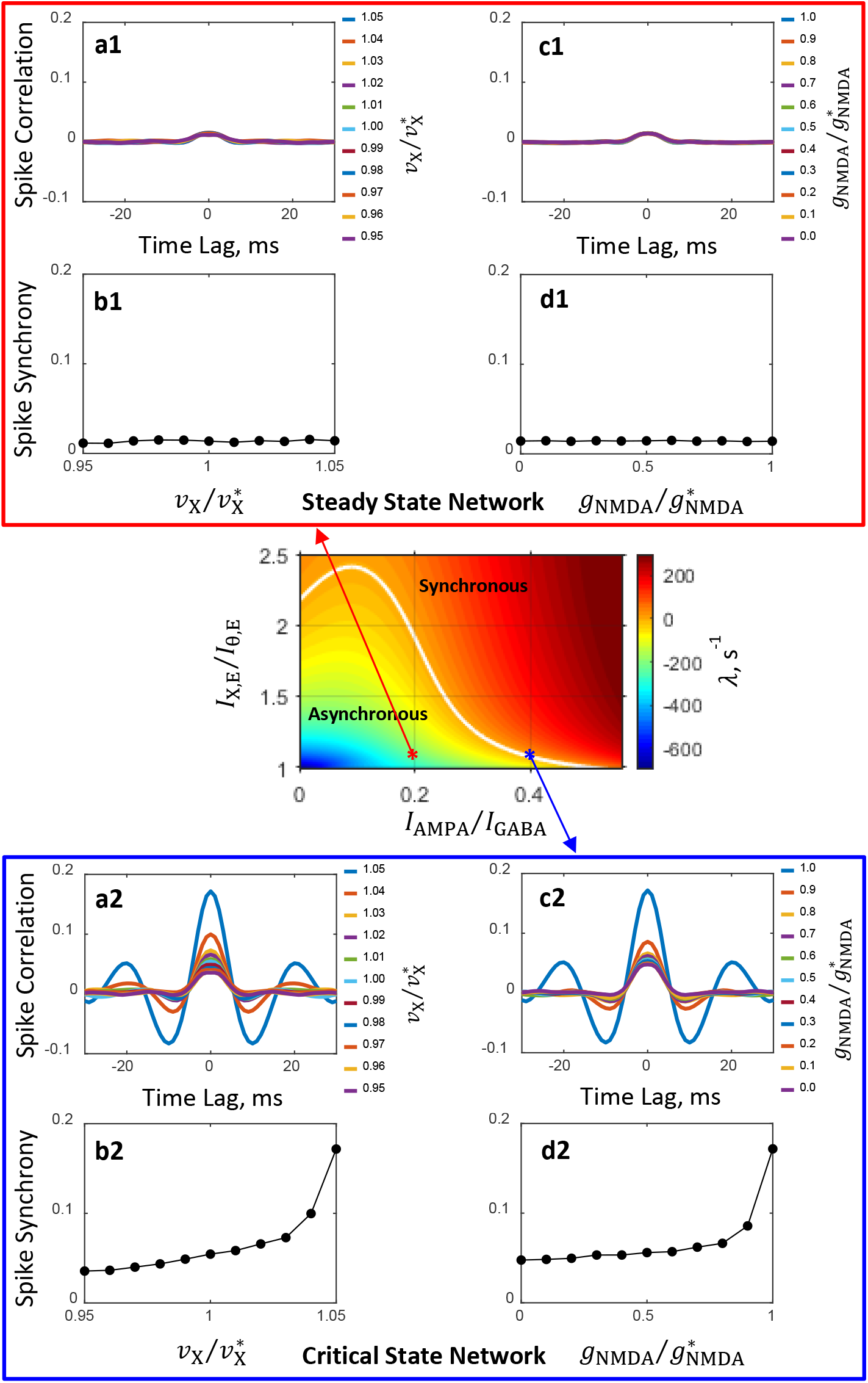
Spiking activity correlation and synchrony computed from spike trains of simulated networks. Conductance parameters are solutions of mean field equations for the steady state primary network (**a1**, **b1, c1**, **d1**) and the critical state primary network (**a2**, **b2**, **c2**, **d2**) corresponding to the red and blue asterisks, respectively, in the (*I*_AMPA_/*I*_GABA_, *I*_X,E_/*I*_*θ*,E_) state plane shown in Fig 2a and inset. For the steady state network correlation and synchrony are weak and insensitive to the modulation of external input spike rate *ν*_X_ (**a1**, **b1**) and NMDAR conductance *g*_NMDA_ (**c1**, **d1**). In contrast, for the critical state network spike correlation depends strongly on the external spike rate (**a2**) and NMDAR conductance (**c2**) and the degree of spike synchrony could be modulated from relatively weak to strong (**b2**, **d2**). Results shown in (**c1**, **d1**, **c2**, **d2**) are obtained from simulations in which *ν*_X_ is increased by 5%. The magnitudes of modulation of *ν*_X_ and *g*_NMDA_ are normalized by their standard values 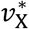 and 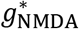, respectively. The numbers next to color-coded markers for spike correlation plots show the normalized magnitudes of external input spike rates, 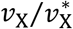, (**a1**, **a2**) and NMDAR conductance, 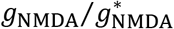, (**c1**, **c2**).

### Circuit mechanisms of spike synchronization modulation

What are the network mechanisms of external drive and NMDA conductance dependent spike synchronization? Why in the network close to the boundary between asynchronous and synchronous regimes, are spike correlations strongly affected by the modulations of external inputs and recurrent NMDA currents, but in the network far from this boundary and deep in the region of the asynchronous regime, correlations are essentially independent of these modulations? How does the interplay between synchronous and asynchronous regimes at their boundary lead to spike synchronization when external input rate *ν*_X_ increases, and to desynchronization when the NMDA conductance *g*_NMDA_ decreases?

To answer these questions and to illuminate the role of asynchronous and synchronous regimes in the shaping of network-wide synchronization of spiking activity, we carried out linear stability analysis in the 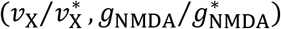 parameter plane while keeping the remaining parameters fixed. For both steady and critical state primary networks, stability is investigated in the vicinity of the standard values of the external input spike rate and NMDAR conductances corresponding to the respective networks.

Figure 5 illustrates state diagrams in the 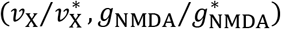 plane in the neighborhood of the steady (Fig. 5a) and critical (Fig. 5b) state primary networks. As in Fig. 2a, the critical line (*λ* = 0) separating the asynchronous stationary (*λ* < 0) and synchronous oscillatory (*λ* > 0) states is shown in white color. Asterisks correspond to the loci of the steady (Fig. 5a) and critical (Fig. 5B) state primary networks in these parameter planes. It is seen that the modulations of *ν*_X_ and *g*_NMDA_ in the steady state primary network (Fig. 5a) do not change the network state; these modulations have no impact on the spike correlation and the strength of synchrony (cf. Fig. 4b1, d1 and Fig. 5a insets).

**Figure 5.**
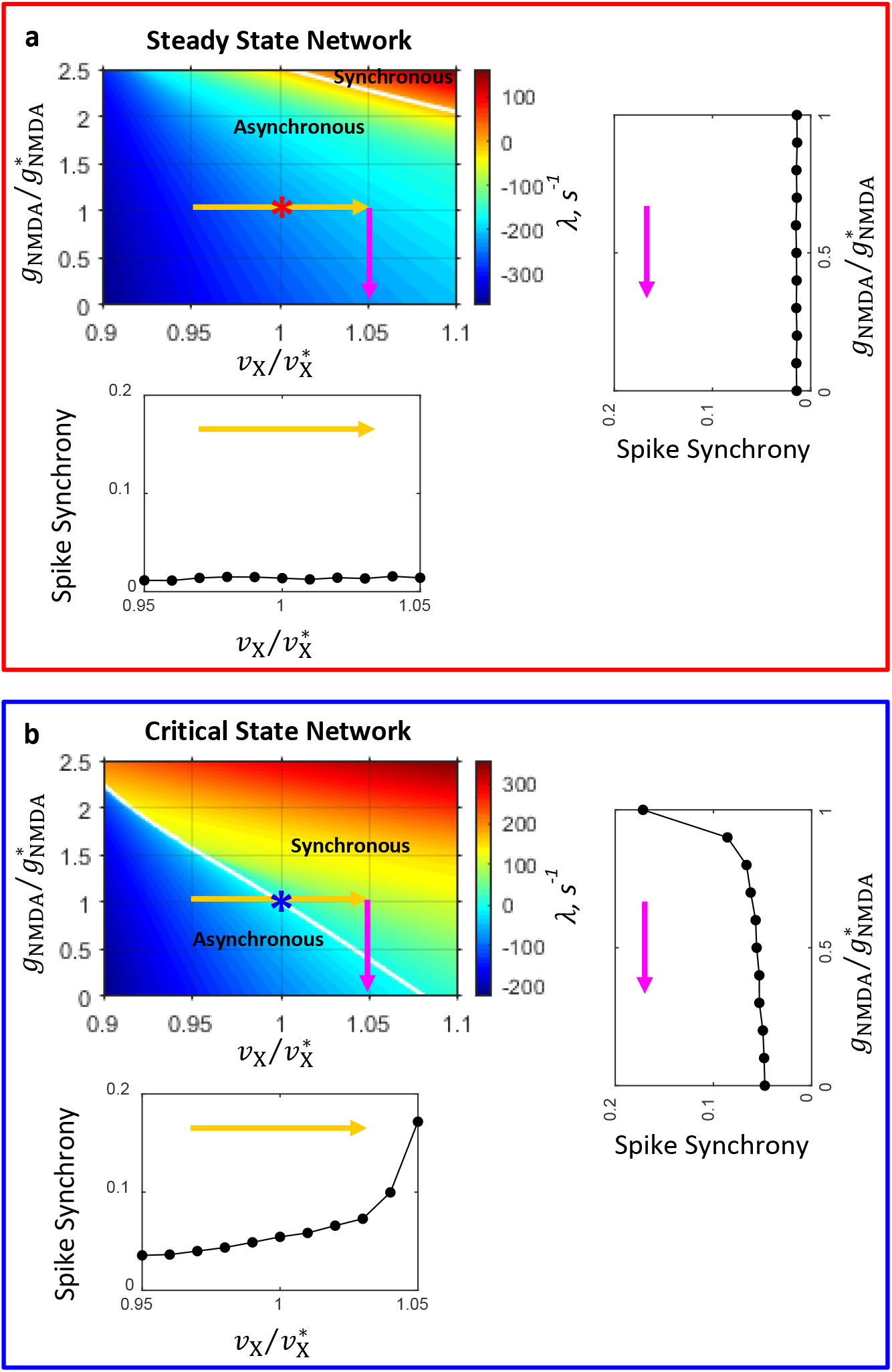
Network state diagrams in the 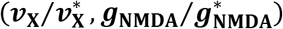 plane. The critical line (*λ* = 0, white line) separates the parameter plane into regions of asynchronous stationary (*λ* < 0) and synchronous oscillatory (λ > 0) regimes. Asterisks correspond to the steady (**a**) and critical (**b**) state primary networks in these planes. Color-coded arrows show the range of modulation of *ν*_X_ (yellow) and *g*_NMDA_ (magenta) corresponding to the range of modulation of these parameters for which temporal correlations of spiking activity and synchrony are shown in Fig. 4. The insets show how spike synchrony changes along the corresponding arrows in the state diagrams. These insets display the same plots for spike synchrony that are shown in panels **b1** and **b2** (bottom insets in **a** and **b**, correspondingly) and **d1** and **d2** (right insets in **a** and **b**, correspondingly) in Fig. 4.

In contrast, the modulations of *ν*_X_ and *g*_NMDA_ in the critical state primary network (Fig. 5b) induce transitions between the network states. Specifically, as the external input spike rate *ν*_X_ increases (horizontal yellow arrow in Fig. 5b) the system crosses the boundary between asynchronous and synchronous regimes and the network state changes from stationary to oscillatory; this transition is accompanied by a sharp increase in spike synchrony (cf. Fig. 4b2 and Fig. 5b bottom inset). The decrease of NMDAR conductance *g*_NMDA_ (vertical magenta arrow in Fig. 5b) causes the system to cross the boundary again, and the network state changes now from oscillatory to stationary; this transition is accompanied by a sharp decrease in spike synchrony (cf. Fig. 4d2 and Fig. 5b right inset).

Thus, this analysis reveals that networks that are close to the boundary between asynchronous and synchronous regimes, in contrast to asynchronous networks that are far from this boundary, have a rich dynamic behavior. The dynamic states of these networks could be easily switched around by modulations in the external drive and the strength of recurrent excitation by NMDAR mediated currents. Switching between the network states, in turn, results in sharp changes in the degree of network-wide synchronization of spiking activity in response to these modulations.

### Explaining the effects of blocking of NMDAR observed in primate PFC by the prefrontal circuit model

As illustrated in Fig. 1b, spiking activity observed in monkey PFC in the DPX task (Zick et al., 2018) remains practically desynchronized after probe presentation for about 200 ms but it begins to increase sharply about 200 ms before the motor response. To get a deeper insight into the properties of spike timing dynamics, we show in Fig. 6 temporal correlations of spiking activity during the 200 ms period following probe presentation (Fig. 6a1) and during the 200 ms period preceding the motor response (Fig. 6b1) in drug-naïve (black) and drug (magenta) conditions. It can be now appreciated that in drug-naïve condition, population activity during the pre-response period develops characteristics of synchronized oscillation behavior, as signaled by the appearance of time lagged peaks of correlation (blue arrows, Fig. 6b1). However, administration of a drug blocking NMDAR desynchronizes neuronal activity during this period (Fig. 6b1).

**Figure 6.**
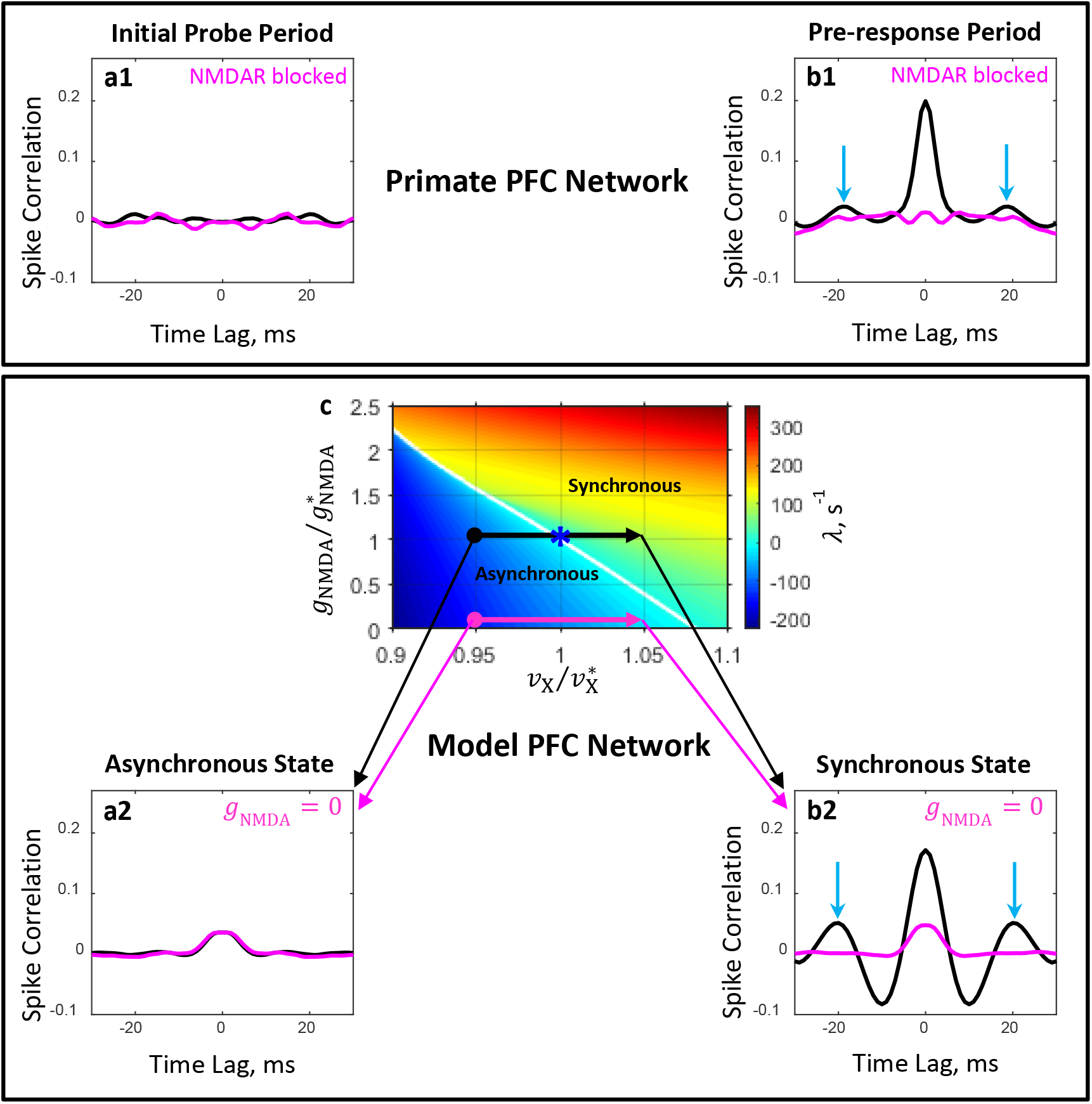
Direct comparison of the effects of blocking of NMDAR in primate PFC and in the prefrontal circuit model. **a1, b1:** Plots show population average temporal correlations between spiking activity of neuron pairs recorded from PFC during the 200ms period immediately following probe presentation (**a1**) and the 200ms period immediately preceding the motor response (**b1**) in the DPX task (Zick et al., 2018). In the drug-naïve condition (black line), population activity during the pre-response period develops characteristics of synchronous oscillation with a frequency of ~55 Hz (peaks at time lags ±18 ms, blue arrows, **b1**). Administration of a drug blocking NMDAR (magenta line) desynchronizes neuronal activity during the pre-response period (**b1**). **a2, b2, c:** Temporal correlations (**a2, b2**) computed from spike trains of simulated networks corresponding to four conditions shown in the 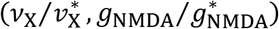 state plane (**c**) by bold dots and arrow heads: *initial probe* (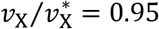, **a2**) and *pre-response* (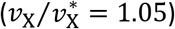, **b2**) periods for *drug-naive* (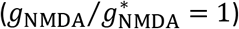, black line) and *drug* (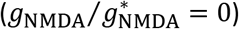, magenta line) conditions. The critical line (*λ* = 0, white line in panel **c**) separates the parameter plane into regions of asynchronous stationary (*λ* < 0) and synchronous oscillation (*λ* > 0) regimes. The locus of the blue asterisk corresponds to the critical state primary network in this plane.

The presence of strong spike synchrony (0 ms lag) together with the correlation peaks at ±18 ms lags in the pre-response period (Fig. 6b1), and the absence of these characteristics in the initial probe period (Fig. 6a1) suggest that after probe presentation but before motor response network dynamics switches from the asynchronous stationary state to the synchronous oscillation state with a *γ*-frequency around 55 Hz. Desynchronization of neuronal activity produced by drug administration implies that NMDAR blockage prevents PFC circuits operating in the asynchronous regime from switching to synchronous dynamics.

These experimental findings could be readily explained by a prefrontal network model that operates on the boundary between asynchronous and synchronous regimes. We start by recalling that in the framework of our approach: 1) the pre-response afferent signals, which we assume are received by PFC neurons before the monkey’s response, are modeled as an increase in the external spike rate from its background level *ν*_X_ and 2) the effect of drug administration is modeled by setting NMDAR conductances *g*_NMDA,E_ and *g*_NMDA,I_ to zero. The capacity of the prefrontal network model to provide a circuit mechanism for the emergence of synchrony in spiking activity and drug-dependent desynchronization can be illustrated by considering the system’s behavior in the 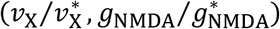 state plane around the point 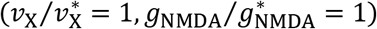 corresponding to the critical state primary network (Fig. 6c). In this space, the effects of probe presentation on the spiking dynamics of the prefrontal circuit model under drug-naïve 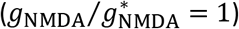 and drug 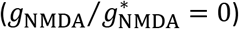 conditions are represented, respectively, by black and magenta horizontal arrows (cf. Fig. 6c). The arrows are pointing from the state of the network corresponding to the initial probe period 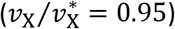 to the network state corresponding to the pre-response period 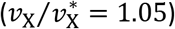.

In drug-naïve condition, increase in the external spike rate *ν*_X_ switches the circuit model from asynchronous to synchronous regime (cf. Fig. 6c, black arrow crosses the boundary between the regimes). The oscillation frequency is about 50 Hz, which is manifested in the temporal correlations of spiking activity as a sharp increase in synchrony and appearance of peaks at ±20 ms lags (cf. Fig. 6 a2 vs b2, black line). This is very similar to what is observed in monkey PFC during the initial probe and pre-response periods in the DPX task (cf. Fig. 6 a1 vs a2 and b1 vs b2, black line).In the drug condition, setting NMDAR conductance to zero prevents the circuit model from switching to the synchronous regime in response to an increase in the external spike rate *ν*_X_ (cf. Fig. 6c, magenta arrow does not cross the boundary between the regimes). This, in turn, considerably reduces the degree of spike synchrony compared to drug-naïve condition (Fig. 6b2, magenta vs black line), similar to the desynchronizing effect of NMDAR antagonist administration on spiking activity in monkey PFC (Fig. 6 b1, magenta vs black line).

In summary, this analysis suggests that because the prefrontal network model operates close to the boundary between asynchronous stationary and synchronous oscillatory regimes it has a considerable capacity to accurately capture experimentally observed aspects of spike synchrony in both drug-naive and drug conditions.

### Role of the balance between NMDAR mediated recurrent excitation and GABA inhibition

So far, in most of our analyses we did not vary the balance between the tonic component of recurrent excitation mediated by NMDA and GABA inhibition, keeping it fixed at 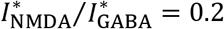. We have only shown that the network frequency at the onset of oscillation essentially is independent of the *I*_AMDA_/*I*_GABA_ balance (Fig. 2–figure supplement 1b), and that the characteristic features of the (*I*_AMPA_/*I*_GABA_, *I*_X,E_/*I*_*θ*,E_) state diagram qualitatively remain unchanged when this balance is varied (Fig. 2–figure supplement 1a). Could, however, the 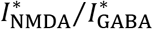 balance be crucial for the prefrontal circuit model capacity to provide the underlying mechanism for external input and NMDA conductance dependent spike synchronization? To investigate this issue, we analyzed how characteristic features of the 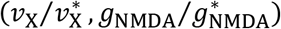 state diagram shown in Fig. 6c depend on the 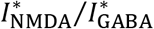 balance.

Figure 7 shows state diagrams in the 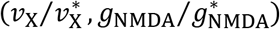 plane obtained for several 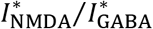 balance values. It is seen that the orientation of the critical line in the state space depends on the 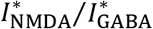 balance. When the balance is shifted toward stronger inhibition (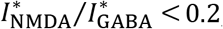, Fig.7a), the critical line becomes too steep: in the drug condition, blocking NMDA current may not necessarily lead to spike desynchronization because the external spike modulation could trigger the network to switch to the synchronous regime (cf. magenta arrow in Fig. 7a). On the other hand, when the balance is shifted toward stronger tonic excitation (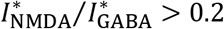, Fig. 7c), the critical line becomes too flat: in the drug-naïve condition the external spike modulation may not be able to produce strong enough synchrony because the system would be too close to the critical line, and not shift deep enough into the region of the synchronous regime (cf. black arrow in Fig. 7c).

**Figure 7.**
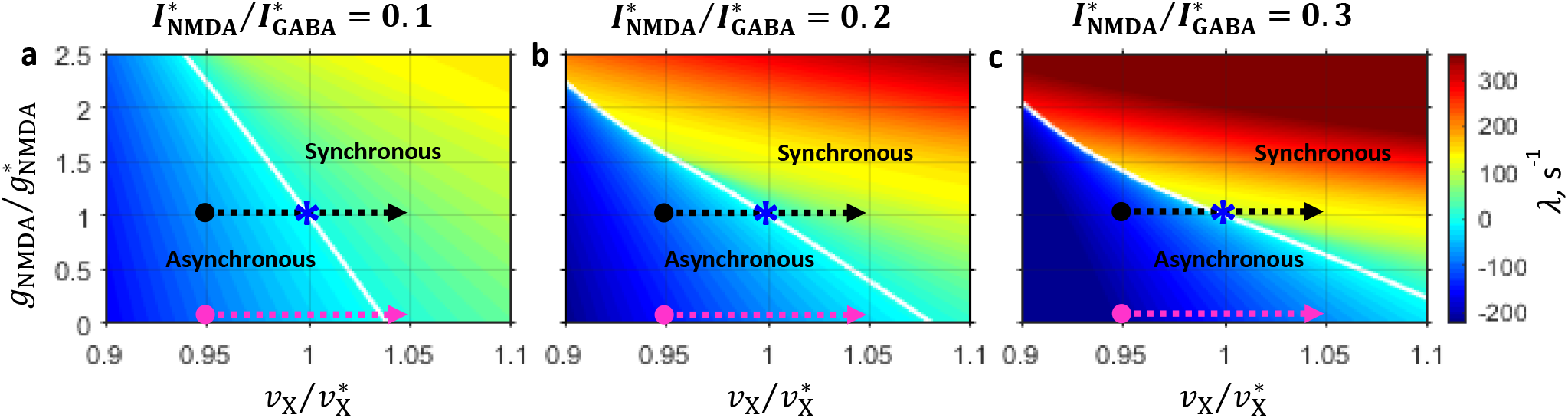
State diagrams in the 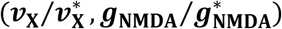 plane obtained for several values of the balance between the NMDA and GABA currents. Notations are the same as in Fig. 6c. **a:** 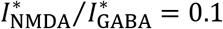; **b:** 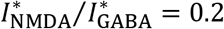; **c:** 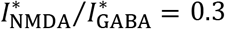. Note that the critical line orientation depends on the 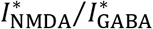 balance.

## Discussion

To investigate how loss of NMDAR synaptic function may distort the physiological dynamics of prefrontal circuits causing those circuits to fail in schizophrenia, and to provide an explanation for biological phenomena we recently observed in monkey prefrontal cortex (Kummerfeld et al., 2020; Zick et al., 2021, 2018), we developed a theoretical framework employing a sparsely connected recurrent network model accounting for AMPAR, NMDAR, and GABAR mediated synaptic currents. We used this platform to investigate how spike timing dynamics in the network depended on the interplay between synaptic conductances, and more specifically, how spike timing was distorted by the loss of NMDAR synaptic function. This allowed us to identify circuit mechanisms through which NMDAR malfunction might reduce spike synchrony in prefrontal circuits, potentially triggering a pathogenic sequence disconnecting these circuits via an activity-dependent process in schizophrenia (Dan and Poo, 2004; Timms et al., 2013; Zick et al., 2018).

Based on the simulation results, we suggest a specific circuit mechanism accounting for the modulation of 0-lag spike synchrony during behavior and its dependence on NMDAR synaptic mechanisms that we have observed in monkey prefrontal cortex (Kummerfeld et al., 2020; Zick et al., 2018). The principal features of this circuit mechanism are as follows: (i) synaptic conductance parameters of the underlying circuit are such that it is in an asynchronous state near a critical boundary in the [NMDAR conductance – external input] parameter plane separating asynchronous and synchronous network states, (ii) small increases in extrinsic inputs push the circuit past this critical boundary into the region of a synchronous state, causing emergence of gamma oscillations in population activity, (iii) 0-lag synchronous spiking between neurons emerges as they stochastically entrain to the gamma population rhythm, (iv) blocking NMDAR currents prevents the circuit from switching to a synchronous regime in response to external inputs, (v) thereby precluding emergence of 0-lag synchronous spiking in neurons. This circuit mechanism offers a reasonable explanation accounting for the task-locked increase in 0-lag spike synchrony that occurs in monkey prefrontal cortex just before the motor response in the cognitive control task (Zick et al., 2018): the upsurge in synchrony could reflect increased synaptic input to prefrontal networks at around this time, potentially from mediodorsal nucleus of thalamus (DeNicola et al., 2020). It also explains why pharmacological blockade of NMDAR attenuates 0-lag spike synchrony before the motor response: the deficit in NMDAR mediated synaptic currents prevents prefrontal networks from switching to a synchronous regime in response to external inputs.

In the circuit model the balance between the AMPA component of recurrent excitation and GABA inhibition controls the network frequency at the onset of oscillation, consistent with results in (Brunel and Wang, 2003). This frequency is virtually independent of the balance between the tonic component of recurrent excitation mediated by the NMDAR and GABA inhibition. However, the balance between the NMDA and GABA currents determines the strength of modulation of the external synaptic input needed for switching between the asynchronous stationary and synchronous oscillatory states in the absence and presence of NMDAR antagonist.

Previous work (Wang, 1999) suggested that NMDAR mediated recurrent excitatory currents have a stabilizing effect on the network activity. Compte and colleagues (Compte et al., 2000) carried out spiking network simulations with different relative contributions of the NMDAR and AMPAR mediated currents to the recurrent excitation and showed that with less NMDA but more AMPA currents, the asynchronous steady state becomes unstable and neurons begin to synchronize, leading to network oscillations in the gamma band. At first glance, these simulation results seem to contradict the experimental findings in (Zick et al., 2018). Indeed, in the neural recording experiments blocking NMDAR caused desynchronization of neurons, whereas in the simulations (Compte et al., 2000) the reduction of NMDAR currents provoked strong synchronization. Our model allows to explain this apparent paradox. In general, the asynchronous state is destabilized and oscillation emerges when an excitatory feedback from the fast AMPA currents becomes sufficiently strong and is followed by a strong inhibitory feedback from the slower GABA currents (Wang, 1999). The excitatory feedback can be enhanced via different mechanisms. In (Compte et al., 2000), it is amplified by increasing the contribution of fast AMPA currents while decreasing the slow NMDA currents to maintain the average level of the excitatory recurrent feedback, thereby leading to network oscillations. In our model, the recurrent positive feedback is amplified by increasing external excitatory currents while the conductance of NMDAR synapses remains unchanged. This induces network oscillation and synchronization of neurons as observed in monkey PFC when NMDAR is not blocked (Zick et al., 2018). However, when the NMDAR conductance is set to zero, the recurrent excitation decreases, so that the increase in external excitatory currents no longer provides a strong enough excitatory feedback. As a result, the network remains in asynchronous state and no increase in synchrony occurs, consistent with observations in (Zick et al., 2018) when NMDAR is blocked.

Our work follows prior modeling studies relating NMDAR synaptic dysfunction to neurodynamical failures in schizophrenia. Most studies are in the context of network models of working memory in prefrontal circuits, and have focused on the destabilization of attractor states (patterns of stable neural activity) during a delay period (when the memory of the stimulus must be retained) (Calvin and Redish, 2021; Compte et al., 2000; Loh et al., 2007; Murray et al., 2014). For example, in the seminal work by Compte and colleagues (Compte et al., 2000) mentioned above, the authors investigated the robustness of working memory storage against external synaptic noise and distraction stimuli in attractor networks. They showed that a concomitant increase of NMDAR and GABAR mediated currents leads to an increase of persistent activity and to a decrease of spontaneous activity, thereby enhancing the resistance of the network to distractors (Brunel and Wang, 2001; Compte et al., 2000). In another prominent work, Murray, Wang and colleagues (Murray et al., 2014), employing an attractor network model, investigated the neural and behavioral effects of synaptic disinhibition induced by the malfunction of NMDAR mediated synapses targeting inhibitory neurons. They demonstrated that disinhibition resulted in a broadening of stimulus selective persistent activity at the neural level, with a concomitant loss of precision, increase in variability over time, and increase in distractibility of stored information at the behavioral level. Finally, a recent study demonstrated that blocking NMDAR reduced gamma frequency oscillations in the membrane potential of neurons in a simulation of auditory cortical networks (Kirli et al., 2014). But in that study, designed to simulate EEG correlates of entrainment of auditory cortex neurons to periodic auditory stimuli, oscillatory activity at the population level was achieved by a network in which individual neurons behaved as oscillators exhibiting highly regular and tightly synchronized spike trains giving rise to synchronously regular network oscillations at the same frequency as the spike firing frequency of individual neurons (Kirli et al., 2014) – a fundamentally different form of network activity than is observed in prefrontal cortex where neurons fire highly irregularly with the frequency that is considerably lower than the frequency of network oscillations, and that we simulate here. Thus, no prior modeling study captures the relationship between NMDAR synaptic mechanisms, spike timing, and network oscillations that we have observed in neural recordings (Kummerfeld et al., 2020; Zick et al., 2018), and for which we provide a theoretical explanation in the current report.

Given that in our neural recording data spiking synchrony peaks around the time of the motor response and the following reward (Zick et al., 2018), and that it is likely to gate synaptic plasticity through Hebbian (Dan and Poo, 2004) and dopamine-dependent (He et al., 2015; Kasai et al., 2021; Yagishita et al., 2014) mechanisms, behavior-timed modulations of circuit dynamics likely play a critical role in structuring synaptic connectivity in prefrontal networks and, therefore, in learning and cognitive function. Correspondingly, given that NMDAR blockage results in the reduction of spiking synchrony on the millisecond time scale (Zick et al., 2018), NMDAR synaptic deficits in schizophrenia would be expected to disconnect prefrontal cortical networks via an activity-dependent process in schizophrenia (Dan and Poo, 2004; Kummerfeld et al., 2020; Uhlhaas and Singer, 2015; Zick et al., 2021, 2018) leading to cognitive impairment. This is a novel pathogenic mechanism with significant implications both for what drives disease and what might slow or reverse it, even after chronic illness. Our simulation results identify a circuit mechanism able to account both for the modulation of 0-lag spike synchrony during behavior and for its dependence on NMDAR synaptic mechanisms. This could provide a platform to investigate how interventions that modulate either network synchrony, 0-lag spike correlation, or their influence on synaptic plasticity may restore functional integrity to prefrontal networks and cognitive function for patients.

## Methods

### Experimental data

For the present theoretical study, we used experimental data obtained in our previous work (Zick et al., 2018). Here, we provide brief descriptions of the experimental task, NMDAR antagonist regimen, and neurophysiological recording methodology employed in that work; details have been reported in (Blackman et al., 2016; Zick et al., 2018).

#### Experimental task

Male rhesus macaque monkeys (8-10 kg) were trained to perform the dot-pattern expectancy (DPX) task. This task is closely related to the AX-CPT (continuous performance task) except that dot patterns replace letters as stimuli. During each trial of the DPX tasks, monkeys maintained gaze fixated on a central target as a cue stimulus (1,000 ms), followed by a delay period (1,000 ms), and a probe stimulus (500 ms) were presented. Monkeys were rewarded for moving a joystick to the left if the cue-probe sequence had been AX (69% of trials), or to the right if any other cue-probe sequence had been presented (AY, BX, BY, collectively 31% of trials). Since the correct response to the X-probe depended on the preceding cue (A or B), the task required both working memory and cognitive control. Both The DPX and AX-CPT measure specific cognitive control impairments in schizophrenia (Barch et al., 2003; Jones et al., 2010).

#### Neurophysiological recording

In our previous study (Zick et al., 2018), we recorded neural activity from the region of the principal sulcus (centered on Brodmann’s areas 46) in the dorsolateral prefrontal cortex of two macaques performing the DPX task. We found that 0-lag synchrony while present in both monkeys was much stronger in one than the other animal. For comparison to spiking dynamics in the present neural network simulation, we used neurophysiological recording data from the monkey that exhibited the strongest 0-lag spike correlation during task performance (Zick et al., 2018). For neurophysiological recording, we used a computer-controlled electrode drive (System Eckhorn, Thomas Recording, GmbH) advancing 16, closely spaced, independently movable glass coated platinum/tungsten microelectrodes into the prefrontal cortex. Electrodes were spaced 400 μm apart, and interelectrode distances in the array spanned 400-1,400 μm. Moving the electrodes in depth and the position of the array within recording chambers over days made it possible to isolate the spiking activity of different neural ensembles, each containing 15-30 individually isolated, simultaneously recorded neurons. The database included in the present study consisted of 47 neural ensembles containing a total of 893 prefrontal neurons. Spike correlation was evaluated within ensembles of simultaneously recorded ensembles using spike trains recorded during DPX task performance (Zick et al., 2018).

#### NMDAR antagonist regimen

We examined the effect of systemic administration of an NMDAR antagonist (phencyclidine, 0.25-0.30 mg/kg IM) on spike timing dynamics in prefrontal local circuits. Neural activity was recorded in a Naïve condition (before first exposure to drug), and a Drug condition (following systemic drug administration) (Zick et al., 2018).

### Spike correlation and synchrony

To estimate correlation between spiking activity of simultaneously recorded neuron pairs as a function of time, we used a similar approach described in (Zick et al., 2018). Correlation is evaluated from spiking activity observed during a time window Δ*T* around a given instant of time *t*. The window size Δ*T*, thus, defines the temporal resolution of time resolved correlation. The interval Δ*T* is subdivided into small time bins of width Δ*t*. Activity of neuron *i* in a given trial at a time bin *t*’ is represented by a binary variable *ξ_i_*(*t*’) that can take on two values: 1 if in the time bin *t*’ one or more spikes are present, and 0 if there are no spikes. Correspondingly, time-lagged joint spike activity of neurons *i* and *j* is described by the product *ξ_i_*(*t*’) × *ξ_j_*(*t*’ + *τ*): it is 1 if neuron *i* fired a spike in the time bin *t*’ and neuron *j* fired a spike in the time bin *t*’ + *τ*; otherwise, it is 0. The duration of the bin Δ*t*, thus, defines the spike timing resolution. We assume that spike firing statistics of neurons do not change during the interval Δ*T*, so that low order moments of the binary variables, such as the mean spike frequencies 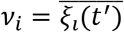 and 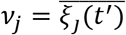 and the mean joint spike frequency 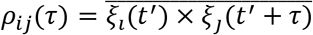, can be reliably estimated by averaging over Δ*T*/Δ*t* time bins (bars 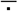 above the expressions denote time averaging operation). To avoid a contribution to correlation from possible cross-trial non-stationarity (slow covariation) of neural activity, for each neuron pair correlation is estimated from single trials and then averaged over all trials. Spiking correlation between neurons *i* and *j* in a single trial is characterized by the observed frequency of joint spikes *ρ_ij_*(*τ*) normalized by the expected joint spike frequency *ν_i_* × *ν_j_* if activity of the neurons were independent: *ρ_ij_*(*τ*)/(*ν_i_* × *ν_j_*). We then average this ratio over the trials to obtain time-lagged correlation of spiking activity as *c_ij_*(*τ*) = 〈*ρ_ij_*(*τ*)/(*ν_i_* × *ν_j_*)〉, where angular brackets 〈·〉 denote trial averaging operation. Finally, *c_ij_*(*τ*) is averaged over the population of simultaneously recorded pairs resulting in the population average spike correlation *C*(*τ*). Spike synchrony is defined as 0-lag correlation.

To accurately estimate spike synchrony and time-lagged correlation in PFC circuits, it is necessary to keep the value of time bin Δ*t*, controlling spike timing resolution, sufficiently small, within 1-2 ms (no more than one spike occurred in a bin). On the other hand, the firing rates of PFC neurons are relatively low, on the order of 10 Hz. Therefore, to increase the number of counts of joint spike events and improve the estimate of spike synchrony while keeping Δ*t* small (and, thus, spike timing resolution sufficiently high), one needs to increase the duration of time window Δ*T* and/or the number of trials *K*. However, Δ*T* should be kept sufficiently short so that during this interval spiking activity remains nearly stationary, whereas *K* cannot be made arbitrarily large because it is limited by practical considerations.

These experimental restrictions, as a result, impose constraints on the firing rates of the neurons in the pair. To derive a meaningful criterion for selecting “good” neuron pairs, we note that for a reliable estimation of the mean joint spike firing frequency, which is a second order statistic, one needs quadratically more experimental samples than for a reliable estimation of the mean spike frequency, which is a first order statistic. We also note that the expected joint spike frequency if neurons in the pair were independent is simply given as the product of their mean spike frequencies. It is this quantity that is used as a reference (normalization) for the quantification of spike correlation strength. Therefore, to reliably estimate the joint spike firing frequency from available samples of a given pair, one should be confident that at least when assuming that neurons fire independently, a sufficiently accurate estimation of the expected joint spike frequency from these samples is possible. This, in turn, means that, given the neuron firing rates *ν_i_* and *ν_j_*, the average total number of counts of joint spikes (*ν_i_*Δ*t*)(*ν_j_*Δ*t*)(Δ*T*/Δ*t*)*K* observed in Δ*T*/Δ*t* bins in *K* trials predicted under the assumption of independence and calculated from experimental samples should be “detectable”, i.e., it should be at least greater than 1. This condition results in a constraint for the geometric mean, 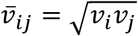, of the firing rates of neuron pairs: 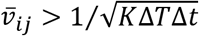. The typical values for the time window and spike resolution are Δ*T*~100 ms and Δ*t*~2 ms. Given that the number of correct trials in the DPX task were on the order of *K*~200, this means that the geometric mean firing rate of neuron pairs, for which a reliable estimation of synchrony can be achieved, should be at least 5 Hz.

### Network model

The network consists of *N* leaky integrate and fire neurons (see, e.g., (Dayan and Abbott, 2001)), of which *N*_E_ = 0.8 *N* are excitatory and *N*_I_ = 0.2*N* are inhibitory (Abeles, 1991; Braitenberg and Schüz, 1998). Neurons are connected randomly with a probability *p*, so that, on average, each neuron receives *C*_E_ = *pN*_E_ connections from excitatory and *C*_I_ = *pN*_I_ from inhibitory neurons. In the framework of mean field consideration, the network is large (*N* ≫ 1) and connections are sparse (*p* ≪ 1) but the average number of connections received by individual neurons, *C*, is large (*C* = *pN* ≫ 1). In most simulations, networks consisted of *N* = 5 · 10^3^ neurons that were randomly connected with the probability *p* = 0.2 and, therefore each neuron, on average, received *C* = 10^3^ connections. In addition, each neuron also receives *C*_X_ external connections from excitatory neurons outside of the network that fire spikes independently according to a Poisson process with rate *ν*_X_.

The dynamics of the membrane potential *V* (*t*) of a neuron below the spike firing potential threshold *θ* obeys the standard leaky integrate and fire equation:

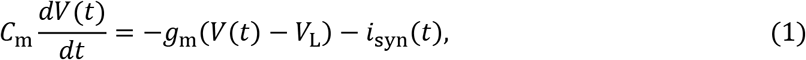

where *C*_m_ is the cell membrane capacitance, is the membrane leak conductance, *V*_L_ is the resting potential, and *I*_syn_(*t*) is the total synaptic current. When the membrane potential reaches the threshold *θ*, the neuron fires a spike, the potential is reset to *V*_rst_, and the neuron becomes insensitive to its input for the duration of a refractory period *τ*_rp_. Both excitatory and inhibitory neurons have *θ* = −50 mV, *V*_L_ = −70 mV, and *V*_rst_ = −55 mV. For excitatory neurons *C*_m_ = 0.5 nF, *g*_m_ = 25 nS, *τ*_rp_ = 2 ms, and for inhibitory neurons *C*_m_ = 0.2 nF, *g*_m_ = 20 nS, *τ*_rp_ = 1 ms (see, e.g., (Koch, 2004)).

The total synaptic input for each neuron is a liner sum of four components:

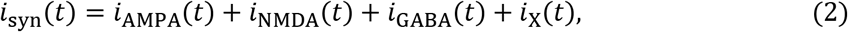

where *i*_AMPA_ and *i*_NMDA_ correspond to recurrent excitatory currents mediated by AMPA and NMDA receptors, respectively, *i*_GABA_ corresponds to inhibitory currents mediated by GABA receptors, and *i*_X_ corresponds to external currents mediated by AMPA receptors. The purpose of external currents is twofold: (i) to represent the noisy inputs due to the background synaptic activity and (ii) to convey neural signals from outside of the network.

The description of component synaptic currents of a postsynaptic neuron follows (Wang, 1999):

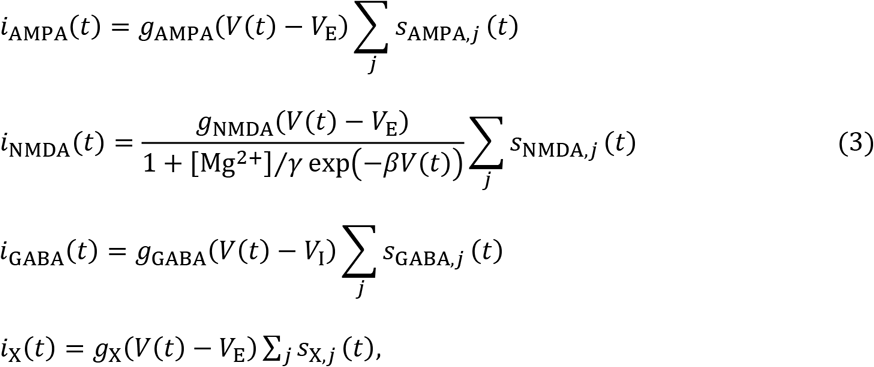

where synaptic reversal potentials *V*_E_ = 0 mV and *V*_I_ = −70 mV. NMDAR mediated currents have voltage dependence controlled by the extracellular magnesium concentration (Jahr and Stevens, 1990): *β* = 0.062 mV^-1^, *γ* = 3.57 mM, [Mg^2+^] = 1 mM. The gating variable *s_R,j_*(*t*), describes the temporal course of postsynaptic currents received from the presynaptic neuron *j* mediated by the receptor *R*, where *R* = X, AMPA, NMDA, GABA. For a spike train generated by a presynaptic neuron with emission themes {*t_k_*}, the temporal dynamics of the gating variable obeys the equations

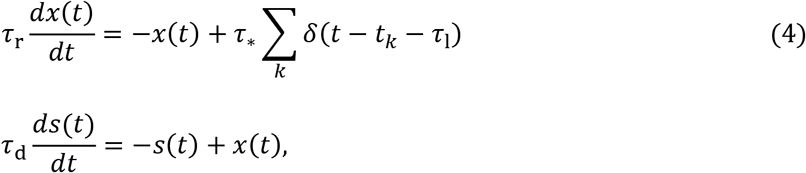

where *τ*_l_, *τ*_r_ and *τ*_d_ are, respectively, latency, rising, and decay time constants. Their values are *τ*_AMPA,l_ = 1 mS, *τ*_AMAP,r_ = 0.2 ms, *τ*_AMPA,d_ = 2 ms for AMPAR mediated currents (Zhou and Hablitz, 1998), *τ*_NMDA,l_ = 1 mS, *τ*_NMDA,r_ = 2 ms, *τ*_NMDA,d_ = 100 ms for NMDAR mediated currents (Hestrin et al., 1990), and *τ*_GABA,l_ = 1 mS, *τ*_GABA,r_ = 0.5 ms, *τ*_GABA,d_ = 5 ms for GABAR mediated currents (Gupta et al., 2000). The time integral of *s*(*t*) in response to a presynaptic spike equals *τ*_*_ and, thus, is independent of the temporal shape of *s*(*t*), which is determined by the rising and decay time constants that are specific to each receptor type. Because the charge flowing to the cell is determined by the product of the time integral of *s*(*t*) and the maximal conductance, we set *τ*_*_ to be the same for all types of receptors, so that the charge entry mediated by each type of receptor is parametrized, in essence, solely by the corresponding maximal conductance parameter.

#### Network simulations

In all direct network simulations, the numerical integration of the coupled differential equations describing the dynamics of membrane potentials and synaptic variables of all cells and synapses were carried out using a custom MATALAB (The MathWorks) code implementing a second order Runge-Kutta method with interpolation of spike firing times between integration time steps Δ*t* (Hansel et al., 1998). In most simulations Δ*t* = 0.1 ms.

#### Mean field approximation

To derive maximal synaptic conductance parameters *g*_X,*α*_, *g*_AMPA,*α*_, *g*_NMDA,*α*_, *g*_GABA,*α*_ (*α* = E,I) providing prescribed neural firing rates *ν*_E_ and *ν*_I_, we used mean field analysis (Amit and Brunel, 1997; Brunel, 2000; van Vreeswijk and Sompolinsky, 1996) extended to networks of neurons with realistic, conductance based synapses (Brunel and Wang, 2001; Renart et al., 2003). For simplicity, we disregard the heterogeneity of synaptic connectivity and assume that each neuron receives *C*_E_ excitatory and *C*_I_ inhibitory connections. In the mean field approximation synaptic inputs are described in terms of their average and their fluctuations arising from both external and recurrent inputs. To this end, the sums of gating variables in Eq 3 are replaced by their respective population averages 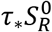, where *R* designates the type of the synapse, and

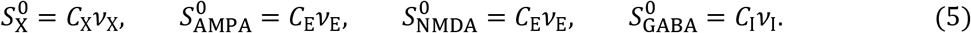

The voltage dependence of NMDAR conductance is linearized around the mean value of the voltage 〈V〉:

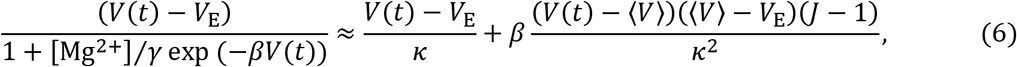

where *κ* = 1 + [Mg^2+^]/*γ* exp (−*β*〈*V*〉). After these simplifications, average components of synaptic currents for excitatory (*α* = E) and inhibitory (*α* = I) populations can be written as

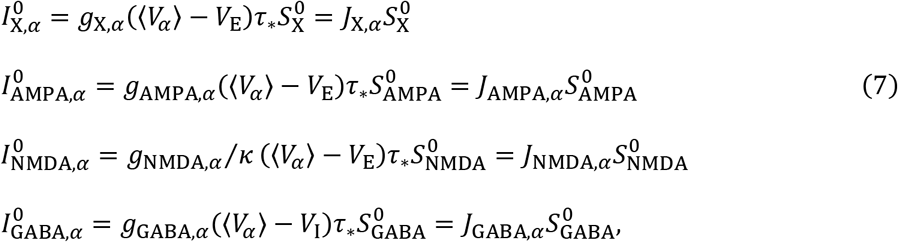

where *J_R,α_* is the effective strength of the *R*-receptor mediated synapse, expressed as the total charge entering the postsynaptic neuron due to a single presynaptic spike. In this framework, the system of equations describing the dynamics of each of individual *N*_E_ excitatory and *N*_I_ inhibitory neurons is simplified to just two equations representing the activity of excitatory, E, and inhibitory, I, populations in terms of the population mean membrane potential *V_α_*(*t*) (Brunel and Wang, 2001; Renart et al., 2003):

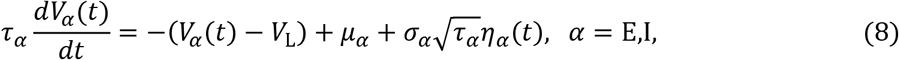

where *V*_L_ is the resting potential, *τ_α_* is the effective membrane time constant, *μ_α_* is the effective mean synaptic input, *σ_α_* is the magnitude of the fluctuations in the synaptic input, and *η_α_*(*t*) is the time course of these fluctuations:

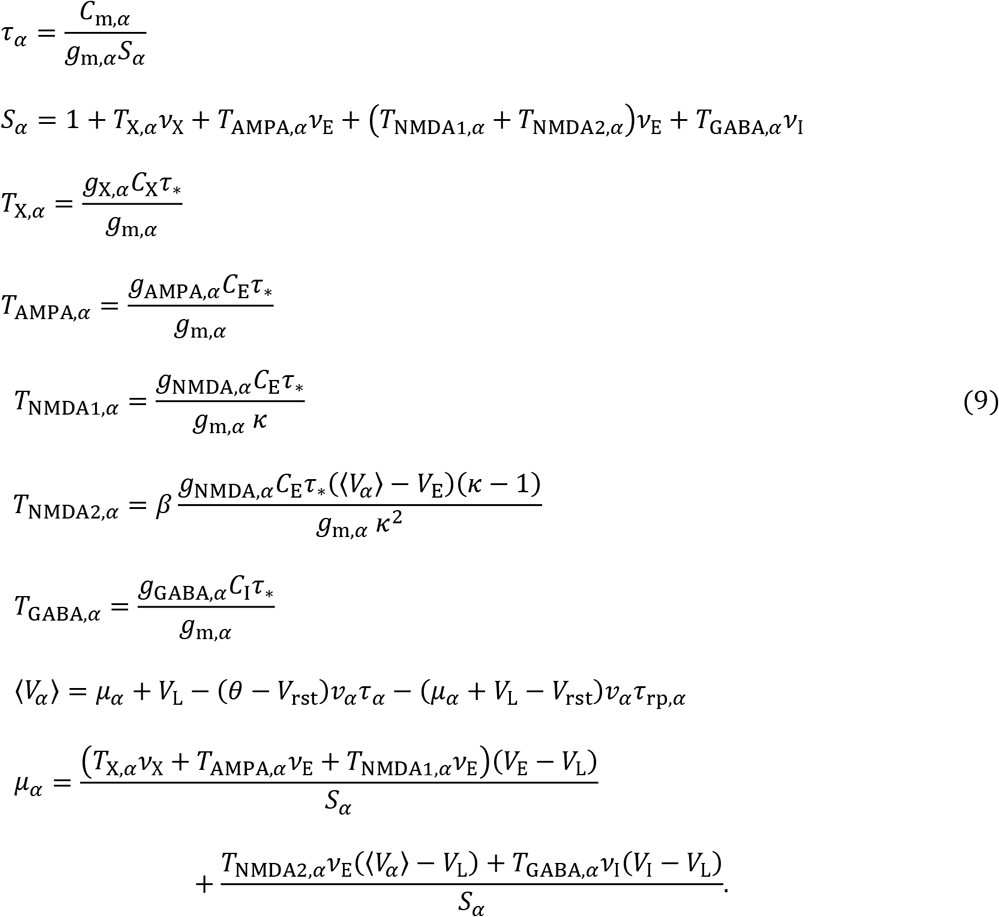

The total synaptic noise 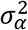 characterizing fluctuations in the input that result from random arrival of spikes is approximated as the sum of the fluctuations in the external and recurrent inputs (Fourcaud and Brunel, 2002):

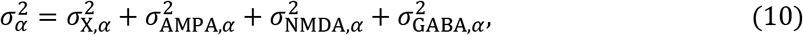

where

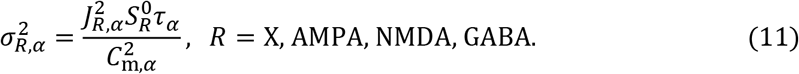

*η_α_*(*t*) is a Gaussian process with zero mean, 〈*η_α_*(*t*)〉 = 0, and an exponentially decaying correlation function, 〈*η_α_*(*t*)*η_α_*(*t*’)〉 ∝ exp(−|*t* – *t*’|/*τ*_syn,*α*_), which is due to synaptic filtering with effective time constant *τ*_syn,*α*_ (Fourcaud and Brunel, 2002):

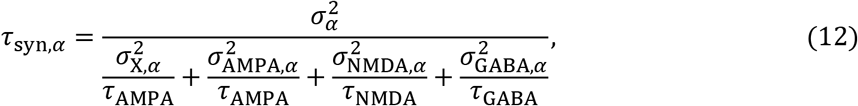

where *τ*_AMPA_ = *τ*_AMPA,l_ + *τ*_AMPA,r_ + *τ*_AMPA,d_, *τ*_NMDA_ = *τ*_NMDA,l_ + *τ*_NMDA,r_ + *τ*_NMDA,d_, *τ*_GABA_ = *τ*_GABA,l_ + *τ*_GABA,r_ + *τ*_GABA,d_ are effective synaptic time constants for AMPAR, NMDAR, and GABAR mediated currents, respectively. In addition, because of sparse connectivity, the correlation of the fluctuations in the synaptic inputs of excitatory and inhibitory populations is neglected: 〈*η*_E_(*t*)*η*_I_(*t*’)) = 0. The firing rate *ν_α_* of a neuron, whose potential is governed by Eq 8, is given by a current-frequency relationship *ϕ*(*μ_α_, σ_α_*) that is a function of the mean and fluctuating part of synaptic input (Brunel and Sergi, 1998; Fourcaud and Brunel, 2002):

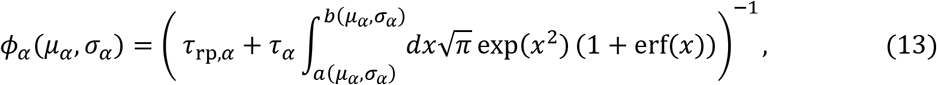

where

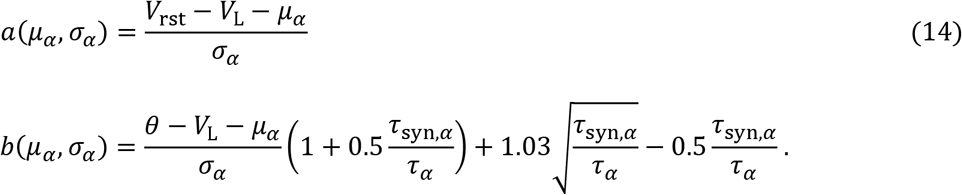

Since *μ_α_* and *σ_α_* themselves depend on the population firing rates *ν*_E_ and *ν*_I_, the two coupled frequency-current equations

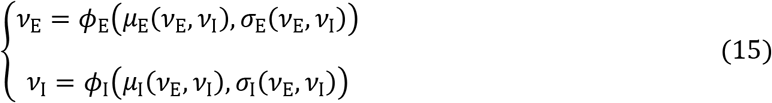

provide a self-consistent description of the network in stationary states, i.e., regimes of network dynamics when the population average quantities such as firing rates and synaptic inputs are constant in time. In the framework of our model, synaptic conductances *g_X,α_*, *g*_AMPA,α_, *g*_NMDA,α_, *g*_GABA,α_ (*α* = E,I) and the external spike rate *ν*_X_ are system parameters controlling the regime of network dynamics; they enter to the mean field analysis through expressions for *μ_α_*, and *σ_α_*,. If these parameters are given, one can solve the self-consistent equations to obtain predicted by the mean field approximation population firing rates 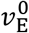 and 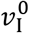 in a stationary state of the network. Conversely, once external *ν*_X_ and population spike rates 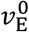 and 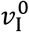 are specified, the self-consistent equations could be solved to find the values of synaptic conductance parameters *g*_X,*α*_, *g*_AMPA,*α*_, *g*_NMDA,*α*_, *g*_GABA,α_ (*α* = E,I) that correspond to these spike rates. However, because there are eight unknown parameters and only two equations, to find a unique solution one would need six additional equations imposing constraints on conductance parameters.

#### Model parametrization

We derive three of these equations by implementing a commonly used constraint (e.g., (Brunel and Wang, 2003; Compte et al., 2000)) that equalizes the ratio of synaptic conductance parameters for component currents in excitatory and inhibitory neurons. Since each component current is proportional to its respective synaptic conductance, this constraint implies that the balance between different components of average synaptic currents 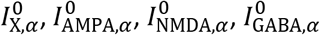 for excitatory (*α* = E) and inhibitory (*α* = I) populations is the same, thus providing the following three equations:

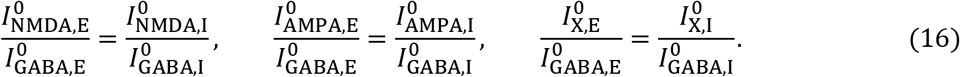

As a result, whenever the ratio of synaptic conductances and/or component currents is involved, the index *α* designating the type of the neuron can be dropped.

Two additional equations are obtained by fixing the balance between inhibition and two-component recurrent excitation at certain values:

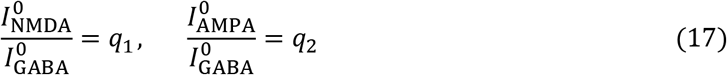

The last constraint is provided in terms of the relative magnitude of average external current of excitatory neurons, 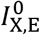:

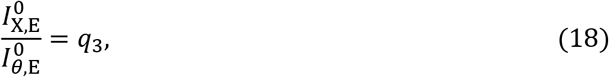

where 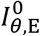 is the current that is needed for an excitatory neuron to reach firing threshold *θ* in absence of recurrent feedback. This approach allowed to parametrize network dynamics in terms of three parameters expressed as ratios of absolute values of average synaptic currents, *I*_AMPA_/*I*_GABA_, *I*_NMDA_/*I*_GABA_, and *I*_X,E_/*I*_*θ*,E_, characterizing the balance between components of recurrent excitation and inhibition, and the balance between external input and firing threshold. For a given external spike rate *ν*_X_ and fixed values of these three parameters, we are now able to solve the self-consistent equations for the eight synaptic conductances that provide the prescribed population firing rates 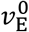 and 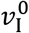 in a stationary state of the network.

We are interested in the asynchronous stationary state in which neurons fire spikes irregularly and at low rates, like neurons in prefrontal cortex. When mean synaptic inputs *μ_α_* are well below threshold *θ*, firing is driven by the synaptic fluctuations *σ_α_* around the mean input, therefore, resulting in irregular spike trains and low rates (Renart et al., 2003). Given that the number of synaptic connections received by individual neurons is large and network connectivity is sparse, solutions of self-consistent equations providing the subthreshold regime for *μ_α_* and, thus, low rate asynchronous network dynamics, arise when inhibition strongly dominates recurrent excitation and the mean external inputs are around or above threshold *θ* (Brunel, 2000; Renart et al., 2003; van Vreeswijk and Sompolinsky, 1996). Thus, for the network to be in asynchronous irregular state the three system parameters characterizing the balance between recurrent excitation and inhibition, and the relative strength of external inputs should be within certain bounds: *I*_AMPA_/*I*_GABA_ + *I*_NMDA_/*I*_GABA_ < 1, and *I*_X,E_/*I*_*λ*,E_ ≳ 1.

#### Linear stability analysis

We perform a linear stability analysis of the asynchronous state (Abbott and van Vreeswijk C, 1993; Brunel and Hakim, 1999)on the basis of an analytical consideration in (Brunel and Wang, 2003). To understand if the network develops instability caused by fluctuations in population firing rates, we consider small deviations from the stationary population rates 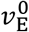 and 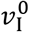. In order to analyze the resulting network behavior, the mean field approach and self-consistent equations providing population mean firing rates 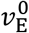 and 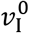 are extended to describe the dynamics of population rates *ν*_E_(*A*) and *ν*_I_(*t*).

In the framework of mean field approximation, each component of synaptic current is determined by the product of effective synaptic strength *J* and average gating variable *S* (cf. Eq 7 for the steady state consideration). The dynamics of *S* is governed by the same type of equations as for the gating variable s of an individual synapse in a given postsynaptic neuron (Eq 4), except that the instantaneous rate of spikes ∑_k_ *δ*(*t* – *t_k_* – *τ*_l_) arriving from the presynaptic cell is replaced by the instantaneous average rate of spikes, *C_α_R__ν_α_R__* (*t* – *τ*_l_), arriving from all presynaptic cells making the same type of synapse in the postsynaptic neuron:

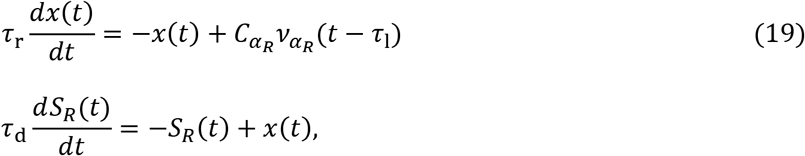

where *R* designates the type of the synapse (*R* = X, AMPA, NMDA, GABA), and *α_R_* designates the presynaptic population establishing these synapses (*α_R_* = X, E for glutamatergic and *α_R_* = I for GABAergic synapse). Since external firing rate *ν*_X_ is stationary, the gating variable for external current is constant in time: *S*_X_ = *C*_X_*ν*_X_. For recurrent currents, the temporal course of *S_R_* is dependent on the instantaneous presynaptic population activity *ν_α_R__*(*t*). Consequently, the total synaptic input current *I*_syn(*t*)_. given as a sum of contributions from external and recurrent components

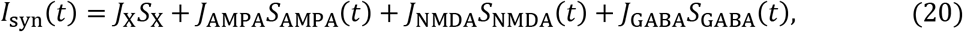

depends on the population firing rates *ν*_E_(*t* and *ν*_I_(*t*). The output firing rate of population neurons, in turn, is determined by the input current and can be modeled in terms of an input-output response function F.

In general, the input-output relationship *ν*(*t*) = *F* (*I*_syn_(*t*)} depends on the spectral characteristics of the input current, resulting in frequency dependent phase shifts and/or amplitude modulations between the oscillatory components of *I*_syn_ and *ν*. However, it has been shown (Brunel et al., 2001; Fourcaud and Brunel, 2002) that the output rate in the leaky integrate and fire neuron model follows instantaneously the temporal variations in its synaptic input current given that synaptic noise is sufficiently strong and synaptic time constant is comparable with membrane time constant. That is, in these conditions, the response does not exhibit a phase shift, and its amplitude is independent of the frequency of oscillatory components of the input current. As a result, even if the input current is varying in time, the input-output function F can be approximated by the current-frequency response function *ϕ*, given by Eq 13, describing the output due to the steady input current.

In the framework of mean field approximation, the output rates *ϕ*_E_(*I*_syn_,E(*t*) and *ϕ*_I_(*I*_syn,I_(*t*)) for excitatory and inhibitory populations must be the same as the instantaneous presynaptic population rates *ν*_E_(*t*) and *ν*_I_(*t*) because both presynaptic and output rates are of the same populations. This requirement results in two self-consistent equations:

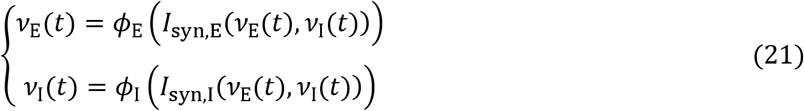

Since the amplitudes of firing rate deviations from the rates in asynchronous steady state are small, *ϕ* (*I*_syn_(*t*) can be linearized about the input current 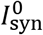 in asynchronous state as:

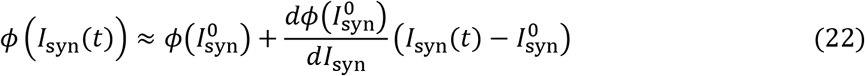

With this approximation, the self-consistent equations for excitatory and inhibitory populations become

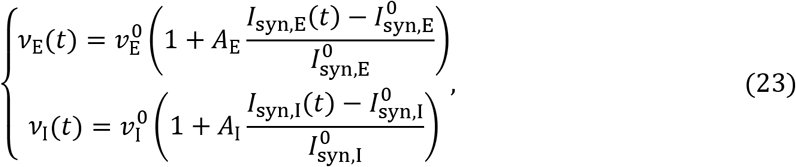

where 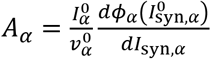 is the dimensionless slope of the current-frequency response function at the current value in asynchronous state, expressed as the ratio between the relative changes in the firing rate and the input current (Brunel and Wang, 2003).

The self-consistent equations Eq 23 together with Eq 19 for the gating variables and Eq 20 for the total synaptic current describe approximate firing rate dynamics of excitatory and inhibitory populations. To determine if the network develops oscillatory instability caused by small fluctuations in population firing rates, we seek solutions for the rates *ν*_E_(*t*) and *ν*_I_(*t*) in which initially small (with relative amplitudes |*ε*_E_| ≪ 1 and |*ε*_I_| ≪ 1) oscillatory perturbations that can change exponentially with time are added to the stationary rates 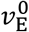 and 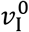 such that: 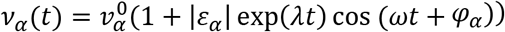 or, equivalently, in complex form

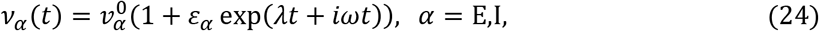

where *λ* is the rate of perturbation growth, *ω* is the oscillation frequency, and *ε_α_* is complex accounting for a possible shift in oscillation phase *φ_α_* between the two populations. We can now replace the firing rates in Eq 19 with these expressions to solve the two equations and determine the synaptic variables *S_R_*(*t*) for recurrent currents mediated by *R* = AMPA, NMDA, GABA receptors:

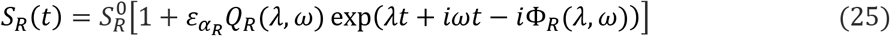

where

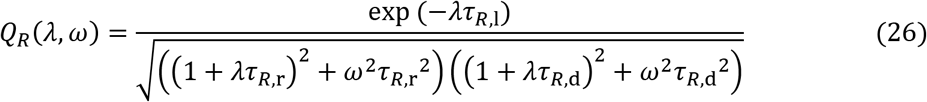

and

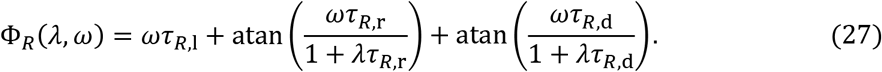

The components of synaptic currents and the total currents *I*_Syn,E_(*t*) and *I*_syn,I_(*t*) can now be calculated and inserted into the linearized self-consistent Eq 23 for population firing rates. Taking into account that the balance between the components of synaptic currents in excitatory and inhibitory populations is equal, we arrive at the following set of two equations:

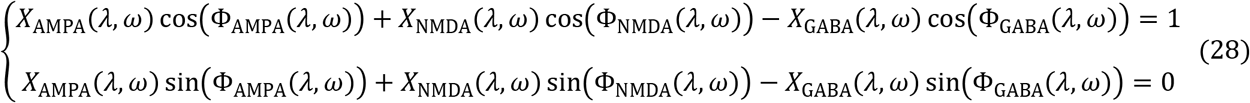

and the relationship between the relative amplitudes:

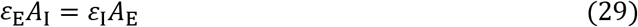

where

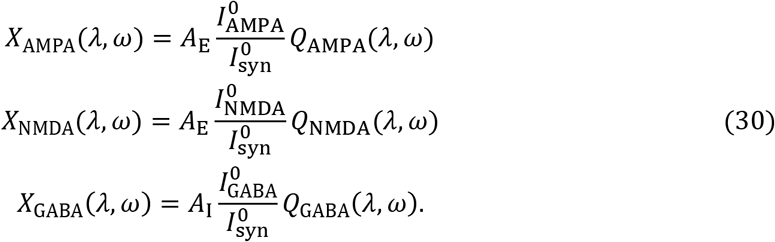

Solving Eq 28, we obtain the rate of perturbation growth *λ* and the oscillation frequency *ω*. Because both *A*_E_ and *A*_I_ are real, the linear relationship between the amplitudes *ε*_E_ and *ε*_I_ given by Eq. 29 means that there is no phase lag between firing rates of excitatory and inhibitory populations.

#### Numerical solutions

Self-consistent mean field equations for the eight conductance parameters, and linear stability equations for the perturbation growth rate *λ* and the oscillation frequency *ω* were both solved numerically using custom codes written in MATALAB (The MathWorks) with the aid of *fsolve* function.

## Data availability

Neural data analyzed in this paper will be uploaded to a public repository (such as Mendeley Data) to be openly accessible upon publication of the paper.

## Code availability

Custom MATLAB codes used for simulations, mean field and stability analysis will be uploaded to a public repository (such as GitHub) to be openly accessible upon publication of the paper.

## Author Information

**David A Crowe**

Department of Biology, Augsburg University, Minneapolis, USA

**Contribution:** Investigation, Formal analysis, Validation, Software, Writing—review and editing

**Competing interests:** no competing interests declared

**Andrew Willow**

Department of Biology, Augsburg University, Minneapolis, USA

**Contribution:** Software

**Competing interests:** no competing interests declared

**Rachael K Blackman**

Department of Neuroscience, University of Minnesota, Minneapolis, USA

**Contribution:** Investigation, Validation, Writing—review and editing

**Competing interests:** no competing interests declared

**Adele L. DeNicola**

Department of Neuroscience, University of Minnesota, Minneapolis, USA

**Contribution:** Investigation

**Competing interests:** no competing interests declared

**Matthew V Chafee**

Department of Neuroscience, University of Minnesota, Minneapolis, USA

**Contribution:** Conceptualization, Investigation, Validation, Writing–original draft, Writing–review and editing, Funding acquisition

**Competing interests:** no competing interests declared

**Bagrat Amirikian**

Department of Neuroscience, University of Minnesota, Minneapolis, USA

**Contribution:** Conceptualization, Methodology, Formal analysis, Software, Visualization, Writing–original draft, Writing–review and editing, Supervision, Project administration

**Competing interests:** no competing interests declared

## Acknowledgments

We would like to provide our special thanks to Apostolos P. Georgopoulos for his intellectual contributions to the study of the temporal correlations in networks and the temporal dynamics of neural ensembles that provided part of the motivation for this work, as well as for his constant support and encouragement. We thank Dean Evans for lab and project management as well as his assistance with surgeries, animal care, and neural recordings; Dale Boeff for his assistance with neurophysiological recording system design and construction, as well as computer programming for signal processing and data analysis; Sofia Sakellaridi for her assistance with neural recordings; Olivia Newman for preliminary analysis. Support for this work was provided by the National Institute of Mental Health (R01MH107491 and P50MH119569 to M.V.C., and R25 MH101076 to R.K.B.), the American Brain Sciences Chair and the Department of Veterans Affairs (to B.A.), the National Institute of General Medical Sciences (T32 GM008244 and T32 HD007151 to R.K.B.), and the Wilfred Wetzel Graduate Fellowship (to R.K.B.). This work was performed while RKB was employed at the University of Minnesota. The opinions expressed in this article are the author’s own and do not reflect the views of the National Institutes of Health, the Department of Health and Human Services, or the United States Government. This material is the result of work supported with resources and the use of facilities at the Minneapolis VA Health Care System. The contents do not represent the views of the U.S. Department of Veterans Affairs, the National Institutes of Health, the Department of Health and Human Services, or the United States Government.

## Ethics

All animal care and experimental procedures conformed to National Institutes of Health guidelines and were approved by the Institutional Animal Care and Use Committees at the University of Minnesota and Minneapolis Veterans Administration Medical Center.

## Figure Supplements

**Figure 2–figure supplement 1.**
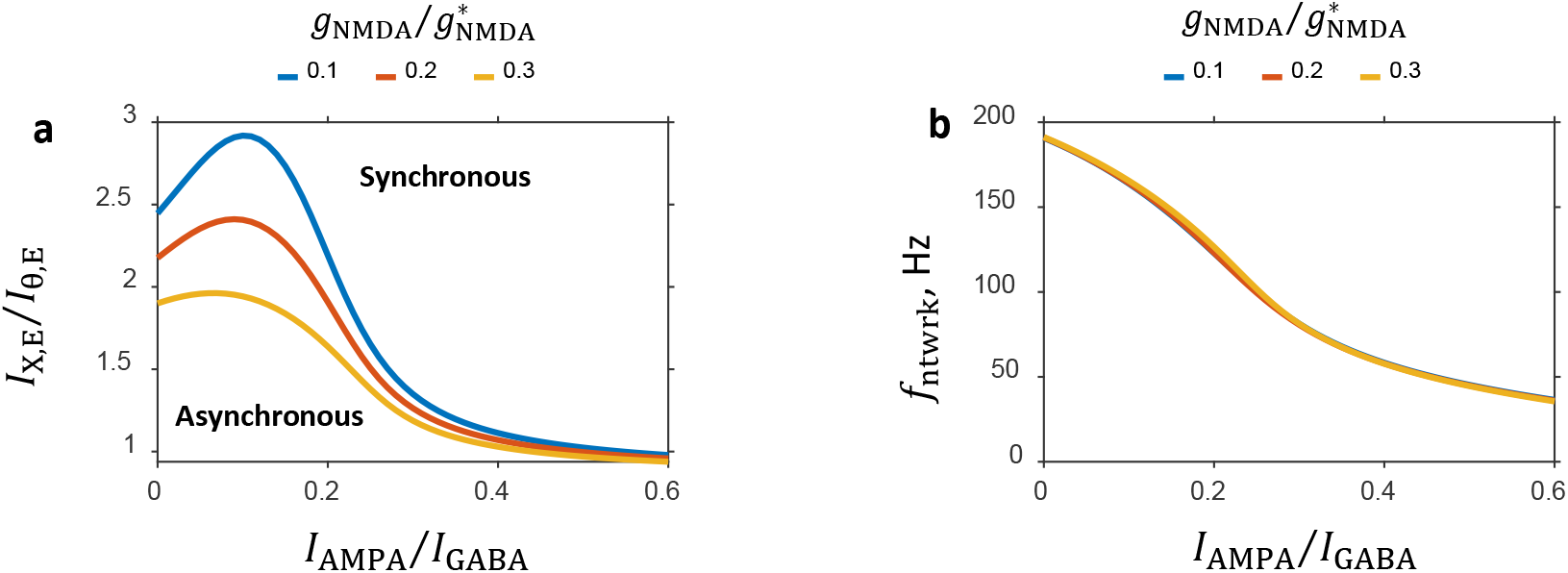
Dependence of the characteristic features of the network on the balance between the NMDA and GABA currents. **a:** Critical line separating the asynchronous and synchronous states in the (*I*_AMPA_/*I*_GABA_, *I*_X,E_/*I*_*θ*,E_) parameter plane is shown for several values of the *I*_NMDA_/*I*_GABA_ balance. **b:** Oscillation frequency on the critical line as a function of the balance between AMPA component of recurrent excitation and GABA inhibition. Plots for different values of the *I*_NMDA_/*I*_GABA_ balance nearly completely overlap.

**Figure 3–figure supplement 1.**
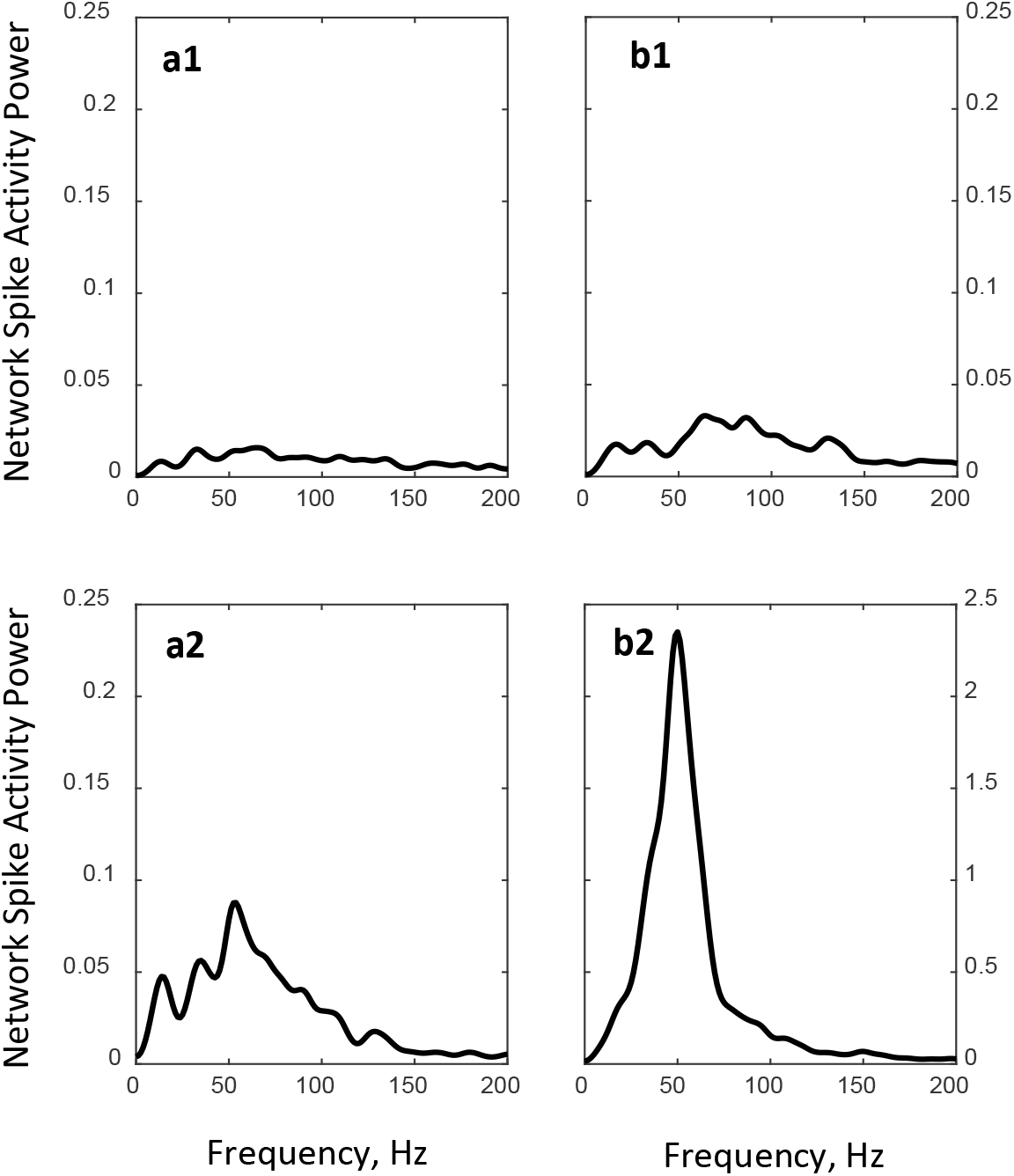
Power spectra of population spiking activity observed in network simulations. **a1, b1:** Spectra of the steady state network activities shown in Fig. 3a1 and Fig. 3b1, respectively. **a2, b2:** Spectra of the critical state network activities shown in Fig. 3a2 and Fig. 3b2, respectively. Note that the scale of Y-axis in **b2** is different from the scale in **a1**, **a2**, **b1**.

